# A microscale platform for the comprehensive analysis of bacterial translation initiation

**DOI:** 10.64898/2026.04.22.719552

**Authors:** Daria S. Vinogradova, Pavel S. Kasatsky, Zoya A. Spiridonova, Sebastian Leyva, Ana Sanchez-Castro, Katherin Peñaranda, Victor Zegarra, Pablo Soriano, Alena Paleskava, Pohl Milon, Andrey L. Konevega

## Abstract

In prokaryotes translation initiation orchestrates protein synthesis through a network of dynamic interactions among the ribosome, mRNA, initiator tRNA^fMet^, and initiation factors (IFs). Traditional approaches that rely on radioactive labeling or surface immobilization are hindered by inherent safety risks and methodological constraints. We present a fluorescence-based analytical platform that integrates microscale thermophoresis (MST) to investigate translation initiation at the molecular level. Employing fluorescently labeled molecules including the initiator tRNA^fMet^, mRNA, and Ifs, enabled a detailed characterization of initiation complex assembly as it progresses from bimolecular to higher-order multicomponent states. To expand the fluorescent toolbox for translation studies we established a novel BODIPY-labeling protocol for 70S ribosomes and confirmed their conformational integrity using nano differential scanning fluorimetry (nanoDSF). Our microscale fluorescent system facilitates probing initiation at a variety of steps, since the role of magnesium ions and initiation factors upon 30S initiation complex formation. The same platform can be applied to investigate the effects of different compounds on translation initiation, as demonstrated for a number of antibiotics, aptamers, and antimicrobial peptides. Using this approach, we determined the antibiotic streptomycin dissociation constant for both 30S and 70S ribosomes, which proved identical at 0.3±0.1 μM, and demonstrated the effect of the antimicrobial peptide rumicidin-1 on translation initiation. Offering a cost-effective and high-sensitivity alternative to conventional methods, this approach advances mechanistic understanding of prokaryotic translation and provides a versatile framework for the discovery of novel protein synthesis inhibitors.

## INTRODUCTION

Messenger RNA (mRNA) translation constitutes a fundamental biological process that decodes genetic information into functional proteins. While the overall mechanism is conserved among archaea, bacteria, and eukaryotes, the initiation phase exhibits substantial divergence across these domains [1]. In bacteria, the rate-limiting step of translation initiation entails coordinated interactions among the 30S ribosomal subunit, mRNA, initiator fMet-tRNA^fMet^ (tRNA^fMet^), and the three initiation factors IF1, IF2 and IF3 [2–4] (Fig. 1). Within the canonical pathway, the stable 30S initiation complex (IC) forms through codon–anticodon pairing between mRNA and tRNA^fMet^, followed by 50S subunit joining to yield the mature 70S IC competent for polypeptide synthesis [4, 5]. This highly coordinated sequence of events governs both the rate and fidelity of translation. The initiation machinery integrates a network of intra- and intermolecular interactions that shape the cellular proteome [4, 6]. Owing to its central role and molecular complexity, the bacterial initiation pathway represents a principal target for diverse translation inhibitors [2, 7, 8].

**FIGURE 1.**
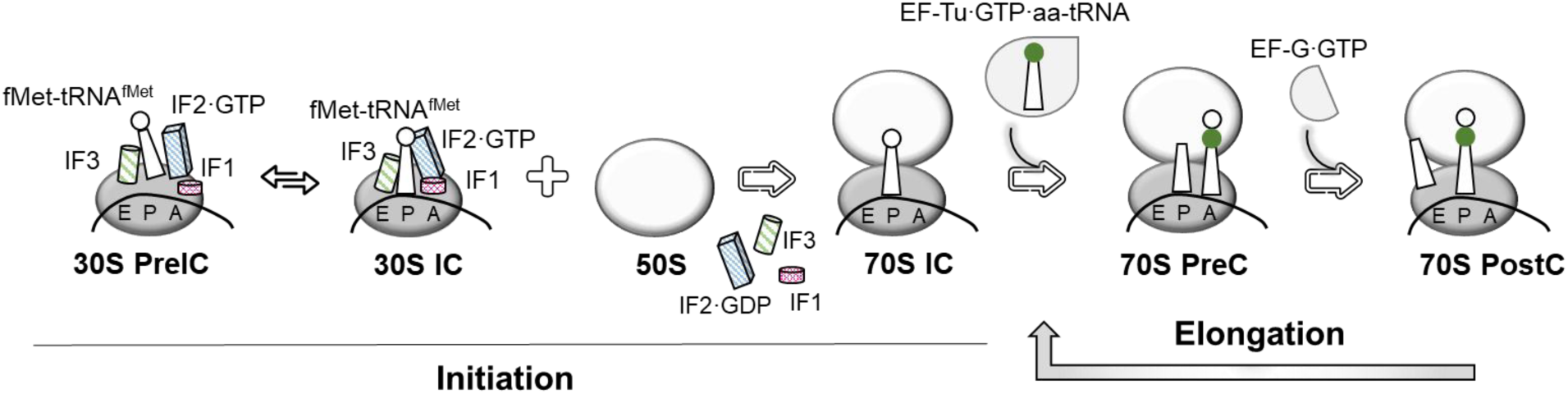
Schematic representation of initiation and elongation bacterial translation cycles. Initiation starts on the binding of the initiator fMet-tRNA^fMet^ and mRNA to the small (30S) ribosomal subunit in the presence of initiation factors (IF1, IF2, IF3), forming a transient 30S pre-initiation complex (30S pre-IC). Upon correct codon–anticodon pairing, the equilibrium shifts toward the assembly of a stable 30S initiation complex (30S IC). Subsequent joining of the large (50S) subunit and release of initiation factors yield the mature 70S initiation complex (70S IC), competent for polypeptide synthesis. The elongation stage, which follows initiation, consists of decoding, peptide bond formation, and translocation steps. The aminoacyl-tRNA (aa-tRNA), as part of a ternary complex with the elongation factor EF-Tu and GTP, is recruited to the ribosomal A site. Following successful codon-anticodon recognition between the aa-tRNA and mRNA, GTP hydrolysis, and dissociation of EF-Tu from the ribosome, peptide bond formation occurs in the peptidyl transferase center of the 50S ribosomal subunit. Following the subsequent translocation of mRNA and tRNA relative to the 30S subunit, catalyzed by the GTPase elongation factor EF-G, the next mRNA codon becomes exposed in the ribosomal A site, ready to accept the cognate aminoacyl-tRNA. The deacylated tRNA leaves the ribosome through the E site. The process ends in the termination stage, where termination factors RF1 or RF2 recognize the stop codon on the mRNA and catalyze the release of the polypeptide chain from the P-site tRNA.

Differences in order, kinetics and affinities of ligands’ interaction with the ribosome reveal the elaborate structure of the bacterial translation initiation process [5, 10–12]. Conventional *in vitro* approaches rely on radioactively labeled ligands. It requires stringent safety procedures and entail substantial disposal costs. Fluorescence-based techniques provide a safer and more versatile alternative for investigating the biophysical and biochemical properties of translation [13–16]. We developed microscale thermophoresis (MST) protocols to investigate translation initiation, an approach that minimizes sample consumption and obviates the use of radioactivity. One of the earliest applications of MST in translation research examined the stringent response in *Escherichia coli* (*E. coli*) by reconstituting the reaction with all essential components, including ribosomes, mRNA, initiator tRNA^fMet^, and initiation factors [15].

Titration experiments using MST enable the determination of binding affinities across a wide range of dissociation constants (10^-12^–10^-3^ M, *K*_D_) and the quantification of inhibition (half-maximal inhibitory concentration, IC_50_) or activation (half-maximal effective concentration, EC_50_) with minimal sample volumes (<5 µL) (Fig. 2a). Fluorescence from ligands—through intrinsic tryptophan residues or covalently attached fluorophores—allows monitoring molecular movement within glass capillaries. An infrared (IR) laser (1480 nm) locally heats approximately 2 nL of sample, generating a temperature gradient of 1–6 K (Fig. 2b). The initial fluorescence measurement serves as a quality control step for detecting aggregation or pipetting inconsistencies. Upon IR laser activation, molecules undergo directed thermal diffusion until an equilibrium is reached; after the laser is switched off, back-diffusion occurs (Fig. 2c). Thermophoretic mobility variations reflect binding-induced changes in molecular conformation, charge, size or hydration shell. Measurements are performed without additional sample immobilization, thereby preserving near-native experimental conditions and highlighting the method’s equilibrium nature [17–19]. The thermophoretic curve reflects fluorescence changes as molecules accumulate or spread away from the observation point during heating cycles (Fig. 2d).

**FIGURE 2.**
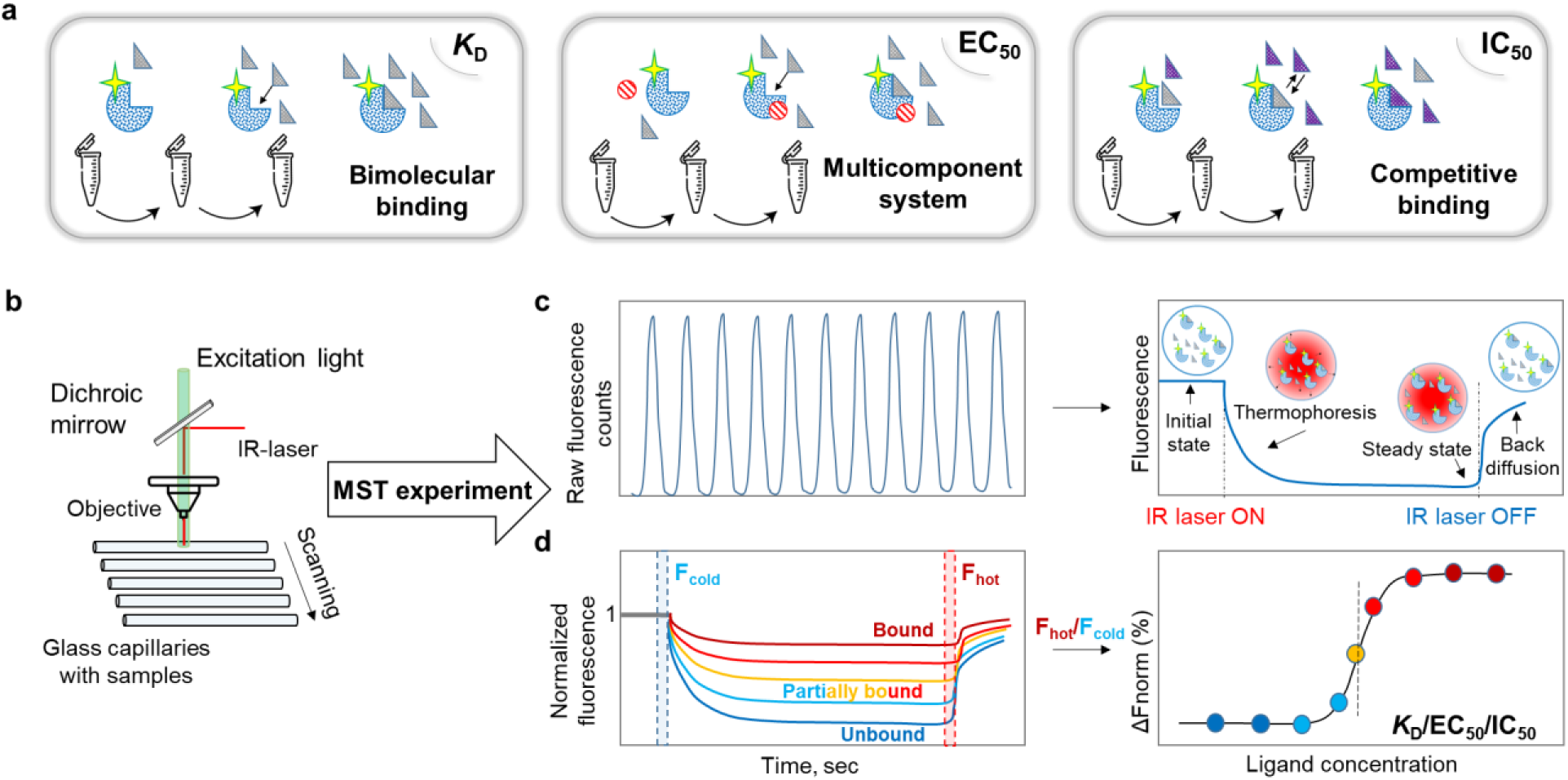
Microscale thermophoresis (MST) technology. (a) Schematic representation of an MST assay used to analyze bimolecular and multicomponent interactions. In a typical titration experiment, a fluorescently labeled reporter molecule (the yellow star symbolizes the fluorophore) is kept at a constant concentration, while the binding partner (ligand) is serially titrated. (b) Reaction samples are loaded into glass capillaries for analysis. (c) The absence of sample aggregation and pipetting errors is verified by monitoring the initial fluorescence signal. Each measurement follows this sequence: recording the initial fluorescence signal ("initial state"), activation of the infrared (IR) laser to induce thermal diffusion ("thermophoresis") until equilibrium ("steady state") is reached, and detection of the reverse diffusion process after the IR laser is switched off ("back diffusion"). (d) Representative MST time traces illustrating ligand binding. The assay monitors one fluorescently labeled ligand at a constant concentration while titrating the unlabeled ligand. Bound (red), partially bound (light red, orange, light blue), and unbound (blue) molecular states display distinct thermophoretic behaviors upon IR heating. The MST response is expressed as the normalized fluorescence ratio during and before heating (F_hot_/F_cold_). The MST signal correlates with the fraction of bound molecules, enabling the determination of ligand affinities and calculation of dissociation constants (*K*_D_), half-maximal effective concentrations (EC_50_), and half-maximal inhibitory concentrations (IC_50_).

We provide detailed quantitative insights into the fundamental steps of bacterial translation initiation using a diverse set of fluorescent reporter ligands. This microscale thermophoresis approach delivers precise binding affinity measurements while offering a cost-effective, accessible platform for investigating core biochemical principles of the process. Notably, the versatility of this approach extends to other complex molecular assemblies, particularly those involving components that are challenging to produce or obtainable only in limited quantities.

## RESULTS

### A Novel Method for BODIPY Labeling of Ribosomes as Reporters

Visualizing ligand interactions on the ribosome is crucial for understanding translation; however, modifications such as fluorescent labeling must be carefully controlled to avoid disrupting molecules function. Methods for labeling ribosomes include modification of purified proteins or RNA followed by reconstitution of subunits [20, 21], the use of reporter oligonucleotides for structural studies [22, 23], the use of site-directed mutagenesis to attach dyes via cysteines [24, 25] or labeling directly at lysine amino acid residues [3]. We introduce a fast and simple method for fluorescently labeling lysine residues in *E. coli* 70S ribosomes with BODIPY (Bpy) dye. Considering the large number of lysine residues in ribosome proteins, we optimized the necessary dye excess to preserve ribosome functional activity while maximizing the sensitivity of fluorescence signal changes during analysis. Importantly, to obtain Bpy-subunits, it is essential to label the entire 70S ribosomes prior to subunit splitting, since labeling the 30S and 50S subunits directly alters the intersubunit surface (Fig. 3a), potentially hampering 70S formation. We assessed the labeling efficiency using SDS-PAGE (Fig. S1a).

**FIGURE 3.**
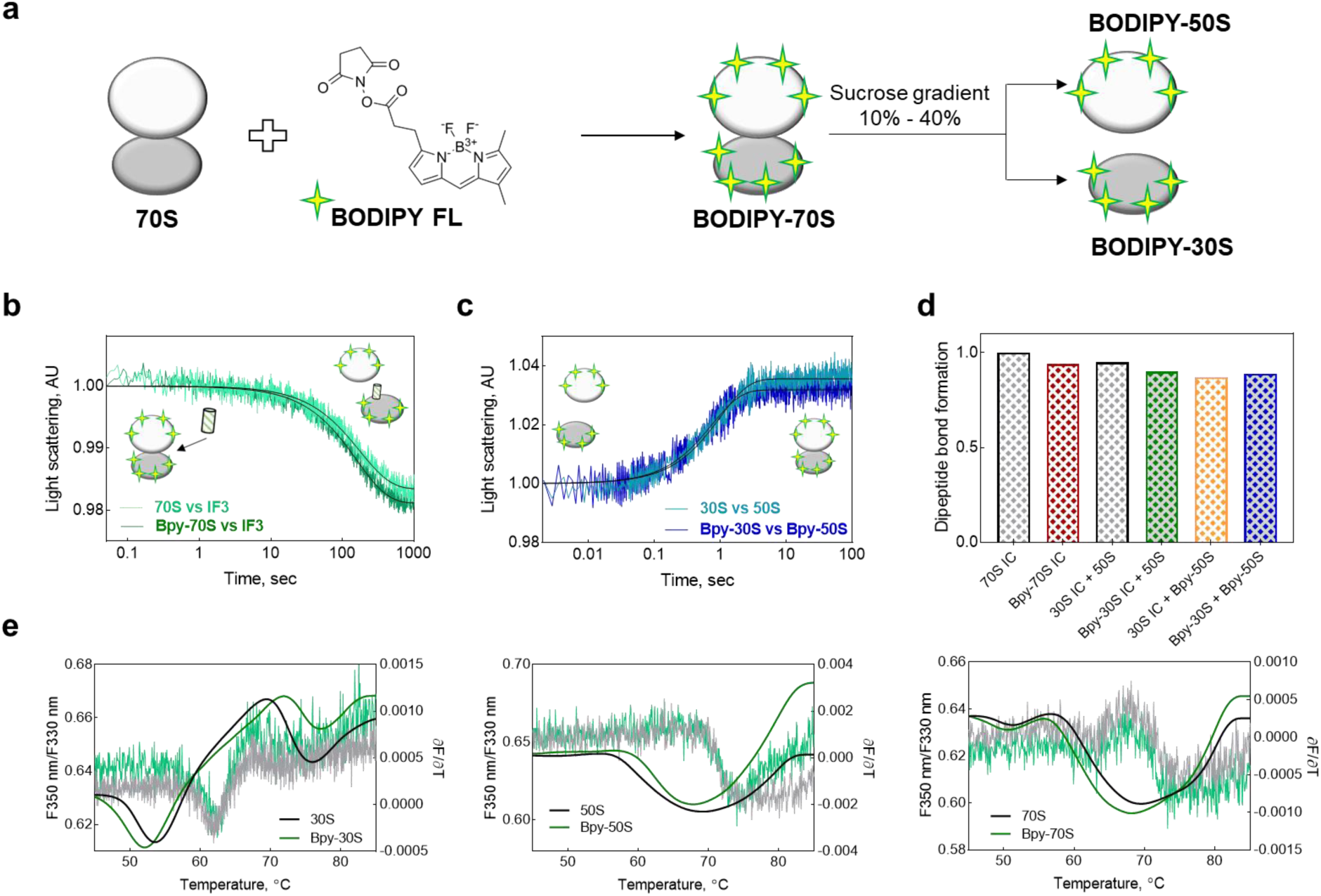
BODIPY fluorescent labeling of ribosomes. (a) Labeling scheme. After dissociation of BODIPY-70S ribosomes, the resulting BODIPY-labeled 30S and BODIPY-50S subunits were separated using a 10%–40% sucrose gradient. (b) Light-scattering analysis of BODIPY-70S ribosome dissociation kinetics in the presence of initiation factor IF3, and (c) re-association of BODIPY-30S and BODIPY-50S subunits compared with native ribosomes. (d) Comparison of dipeptide synthesis efficiency using native and BODIPY-labeled 70S, 30S, and 50S ribosomes. (e) Melting curves of native and fluorescently labeled 30S, 50S, and 70S ribosomes measured by nanodifferential scanning fluorimetry (nanoDSF).

### Functional activity of BODIPY-ribosomes

The association of 30S and 50S subunits into the 70S ribosome as well as the reverse process of dissociation into subunits are critically important for maintaining a pool of functional ribosomes by regulating the balance between monomers and dimers [31–32]. We measured the kinetics of both reactions by stopped-flow technique monitoring the light-scattering signal change. The association rates of native and fluorescently labeled 30S and 50S subunits were very close, yielding values of 1.01 ± 0.02 s^-1^ and 1.33 ± 0.04 s^-1^, respectively (Fig. 3c). The kinetics of 70S dissociation into subunits was measured in the presence of IF3 to exclude the re-association step [26–30]. The dissociation rates of native ((5.6 ± 0.1) x 10^-3^ s^-1^) and Bpy-70S ((6.2 ± 0.1) x 10^-3^ s^-1^) ribosomes were also virtually indistinguishable (Fig. 3b). Importantly, the amplitudes of all the corresponding light-scattering signals were essentially identical, highlighting the equal amount of the functional ribosomes. Thus, fluorescent labeling of ribosomes had no impact on the structural integrity of the intersubunit interface and the dynamics of the ribosome assembly — critical hallmarks of translation efficiency.

Although the fluorescent dye uniformly localizes to ribosomal subunit surfaces without making significant contributions to the functionally active centers, we analyzed signal variations under translation initiation kinetics conditions. We monitored the formation of the 30S IC and 70S IC by recording changes in the fluorescence of labeled ribosomes (Fig. S1b). As expected, assembly of the 30S initiation complex elicited no noticeable fluorescence signal shift, as ligand binding occurs in the intersubunit interface, unlabeled region. Use of Bpy-70S or Bpy-30S and Bpy-50S ribosomes resulted in a marked increase in the fluorescence signal upon 70S initiation complex formation, with observed rates of 0.075±0.004 min^-1^ and 0.049±0.006 min^-1^ (Fig. S1b), respectively. To confirm the functionality of the fluorescent initiation complexes, we tested their ability to participate in peptidyl transferase reaction (Fig. 3d). The use of Bpy-labeled 70S ribosomes, as well as isolated Bpy-30S and Bpy-50S subunits, supported efficient peptide bond formation comparable to that of native ribosomes.

### Probing Ribosomal Stability with Nano Differential Scanning Fluorimetry

Differential scanning calorimetry (DSC) and nano differential scanning fluorimetry (nanoDSF) are classical approaches for assessing the conformational stability of protein preparations [33–36]. We applied nanoDSF technology to monitor the preservation of conformational stability of native and fluorescently labeled 70S ribosomes, as well as in their 30S and 50S subunits (Fig. 3e). Interestingly, 30S subunit and 70S ribosome showed lower thermal stability (melting temperatures – Tm – 53.6°C and 51.3°C, respectively) than the 50S subunit, which remained stable up to 70.1°C under conditions of the experiment. The higher stability of the large ribosomal subunit is most likely based on the high content of rigidly ordered and densely packed rRNA, forming among other things the evolutionarily most ancient parts of the ribosome, such as the peptidyl transferase center and the nascent peptide exit tunnel. The melting profiles of fluorescently labeled ribosomes were indistinguishable from those of their native counterparts (Fig. 3e), confirming that labeling did not disrupt ribosomal stability. To more thoroughly examine the impact of the performed modification on ribosome conformational stability, we conducted nanoDSF analysis under varying ionic conditions, which play a decisive role in ribosomal structural and functional organization [37–43]. Among the major contributors, magnesium (Mg^2+^) and potassium (K^+^) ions are particularly essential to maintain the stability and translational competence of the 70S ribosome. Mg^2+^ primarily governs the structural integrity of the ribosome, whereas K^+^ ions influence both stability and translational dynamics [42, 44–51] (Fig. S1c). Melting curves of native and BODIPY-labeled 70S ribosomes, as well as 30S and 50S ribosomal subunits, remained identical across varying ionic conditions. Only the 50S subunit exhibited reduced sensitivity to Mg^2+^ concentration changes, which did not affect its activity in our conducted control analyses. A more detailed analysis of conformational stability dependence for native and modified ribosomes across varying ionic conditions is available in the Supplementary Materials.

### Bimolecular Ligand Interactions in Translation Initiation Probed by MST

Assembly of the bacterial translation initiation complex proceeds through the multiple sequential stages, requiring the presence of all ligands for optimal efficiency. Individual components, including IFs, tRNA^fMet^, and mRNA form bimolecular complexes with the 30S ribosomal subunit with varying affinities [4, 11, 60–62]. We initiated MST-studies of translation initiation through step-wise analysis. Initially, we examined bimolecular interaction affinities by determining dissociation constant (*K*_D_) values of ligands. We quantified the binding affinities of the IF1, IF2, IF3, mRNA, and tRNA^fMet^ to the 30S subunit, employing each as a fluorescent reporter in MST assays.

IF1, IF2, and IF3 were site-specifically fluorescently labeled with Cy5 via maleimide chemistry, targeting solvent-accessible cysteine residues to yield IF1^Cy5^, IF2^Cy5^, and IF3^Cy5^ (Fig. S2a, b). The measured dissociation constants were 214 ± 98 nM for IF1, 73±32 nM for IF2, and 9±8 nM for IF3 in excellent agreement with values obtained previously by pre-steady-state kinetics (Fig. 4a, b, S2c) [10].

**FIGURE 4.**
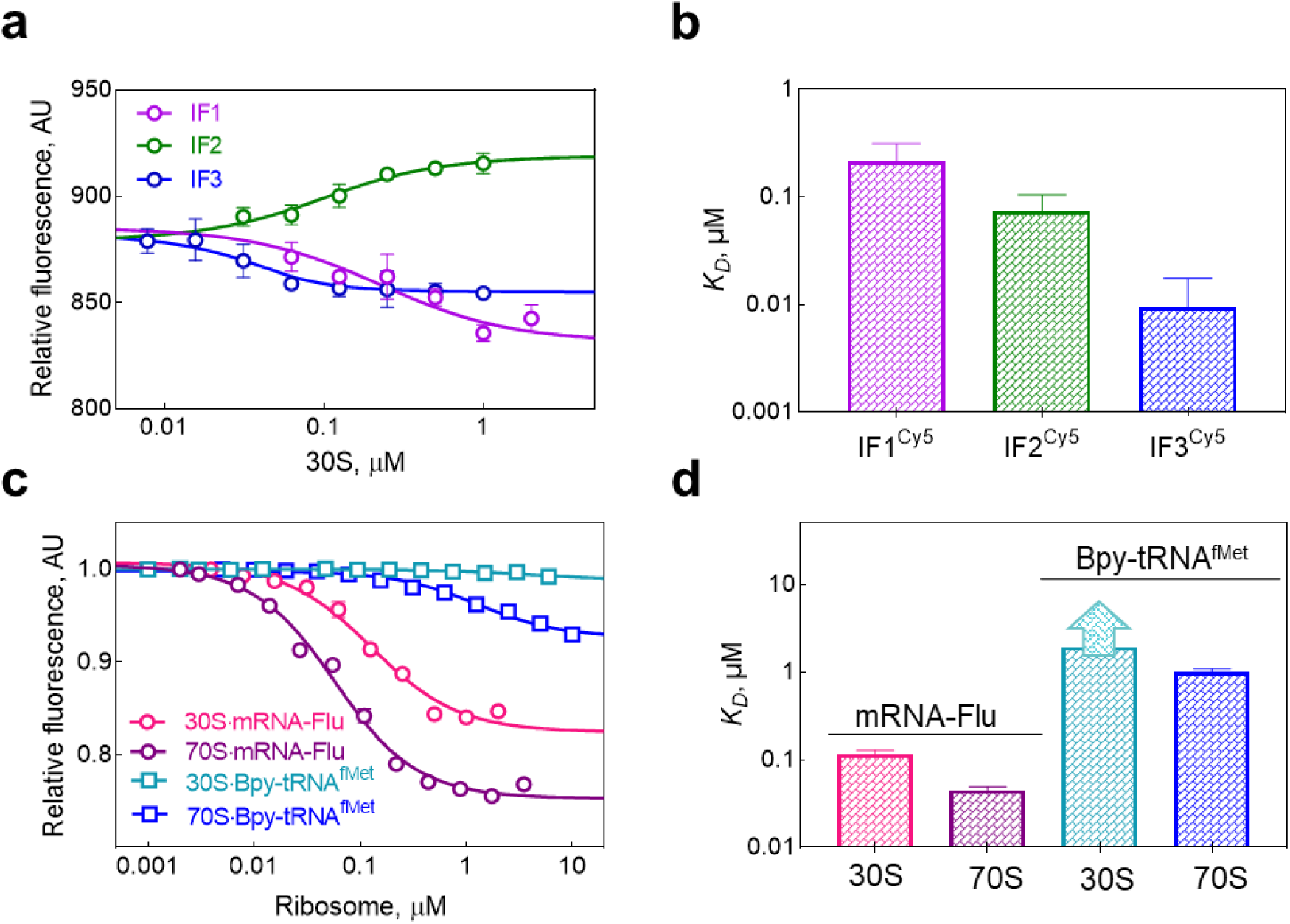
Determination of dissociation constants (*K*_D_) using different fluorescently labeled ligands as reporters. (a) Binding curves of IF1^Cy5^, IF2^Cy5^, and IF3^Cy5^ to the 30S ribosomal subunit. (b) The *K*_D_ values of IF1^Cy5^•30S (magenta), IF2^Cy5^•30S (green), and IF3^Cy5^•30S (blue) complexes. (c) Binding curves of mRNA-Flu or Bpy-tRNA^fMet^ to the 30S or 70S ribosomes. (d) The *K*_D_ values of mRNA-Flu•30S (magenta), mRNA-Flu•70S (violet), Bpy-tRNA^fMet^•30S (light blue), Bpy-tRNA^fMet^•70S (blue) complexes. Error bars represent standard deviations from three independent measurements.

Although mRNA and tRNA^fMet^ can bind the 30S subunit independently, the 30S•mRNA complex exhibits substantially greater stability than the 30S•tRNA^fMet^ complex [63–66]. 30S•mRNA association is mediated by complementary base pairing between the anti-Shine–Dalgarno (anti-SD) sequence in 16S rRNA and the Shine–Dalgarno (SD) sequence in mRNA, positioning the mRNA within a dedicated channel on the subunit surface [4]. In contrast, formation of the 30S•tRNA^fMet^ bimolecular complex is less stable due to the requirement for the presence of IF2, which is responsible for recruiting tRNA^fMet^ to the P site of the ribosome [11, 67].

We quantified the bimolecular interactions of mRNA and tRNA^fMet^ with 30S and 70S ribosomes using MST with fluorescein-labeled mRNA (mRNA-Flu) or BODIPY-labeled tRNA^fMet^ (Bpy-tRNA^fMet^) as the reporters (Fig. 4c, d, S2d). mRNA-Flu exhibited high-affinity binding to 30S and 70S, with dissociation constants (*K*_D_) of 115 ± 14 nM and 45 ± 4 nM respectively. Bpy-tRNA^fMet^ bound efficiently to 70S ribosomes (*K*_D_ = 1.0 ± 0.1 μM), but showed markedly weaker affinity for 30S alone (the *K*_D_ value could not determined).

### MST Measurements of Initiation Complex Formation Efficiency

Bacterial translation initiation represents a multifaceted process that can proceed via distinct pathways involving either 30S or 70S ribosomes [4, 60, 68–71]. In the canonical pathway, formation of the 30S initiation complex is orchestrated by synergistic interactions of IF1, IF2, and IF3, which modulate mutual binding affinities to optimize tRNA^fMet^ recruitment to 30S [4, 10]. Codon–anticodon recognition stabilizes the 30S IC, enabling subsequent 50S subunit joining [4, 72], IF dissociation, and progression to the elongation-competent 70S IC [4, 9, 10, 73–79]. Non-canonical 70S initiation complex formation pathways include leaderless mRNA initiation [70–71], translation re-initiation [80, 81], polysome-associated initiation [82–84], and 70S scanning [69].

We progressively expanded the system toward a multicomponent complex using Bpy-tRNA^fMet^ as the reporter molecule. Elevated concentrations of Mg^2+^ stabilize the tertiary structure of rRNA, potentially facilitating binding of mRNA and tRNA^fMet^ to the 30S subunit [42, 52, 63, 66]. Thus, we examined initiator tRNA^fMet^ binding to the 30S•mRNA bimolecular complex (Fig. 4c) across a range of magnesium ion concentrations (Fig. S3a). We observed robust ternary 30S•mRNA•Bpy-tRNA^fMet^ complex formation under increasing Mg^2+^ conditions (3.1–100 mM) (Fig. S3a). Subsequent system expansion involved adding individual initiation factors to dissect their roles, with each titrated against the 30S•mRNA•Bpy-tRNA^fMet^ ternary complex to assess contributions to tRNA binding efficiency. Considering the role of magnesium ions in Bpy-tRNA^fMet^ binding to the ribosome (Fig. S3a), we conducted experiments at 7 mM or 20 mM Mg^2+^ (Fig. S3b). Thermophoretic shifts were observed exclusively with IF2, producing the expected signal consistent with established data on its key role in recruiting tRNA^fMet^, particularly by accelerating the kinetics of its association [85–87]. Additionally, we demonstrated the versatility of the system by analyzing 30S IC formation efficiency as a function of IF3 in the presence of mRNA lacking a canonical start codon and regulatory elements (polyU mRNA) (Fig. S3c). 30S ICs were practically not formed in the presence of polyU mRNA, potentially reflecting both the absence of productive interaction and the proofreading function of IF3 during codon-anticodon checking.

We next investigated 30S IC and 70S IC formation as a function of mRNA, using initiator Bpy-tRNA^fMet^ or Bpy-70S as reporter molecules or performing label-free measurements, while sequentially adding initiation factors to the system (Fig. 5). mRNA titration revealed maximal efficiency of 30S IC formation when all three IFs were present (Fig. 5a). In the absence of IF3 (or IF3+IF1), mRNA affinity dropped ≈2-fold, accompanied by a weaker thermophoretic signal. Absence of IF1 alone caused a ≈3-fold affinity decrease without substantially affecting signal amplitude (Fig. 5a, d). These findings demonstrate the synergistic contributions of IF1, IF2, and IF3 to 30S IC formation.

**FIGURE 5.**
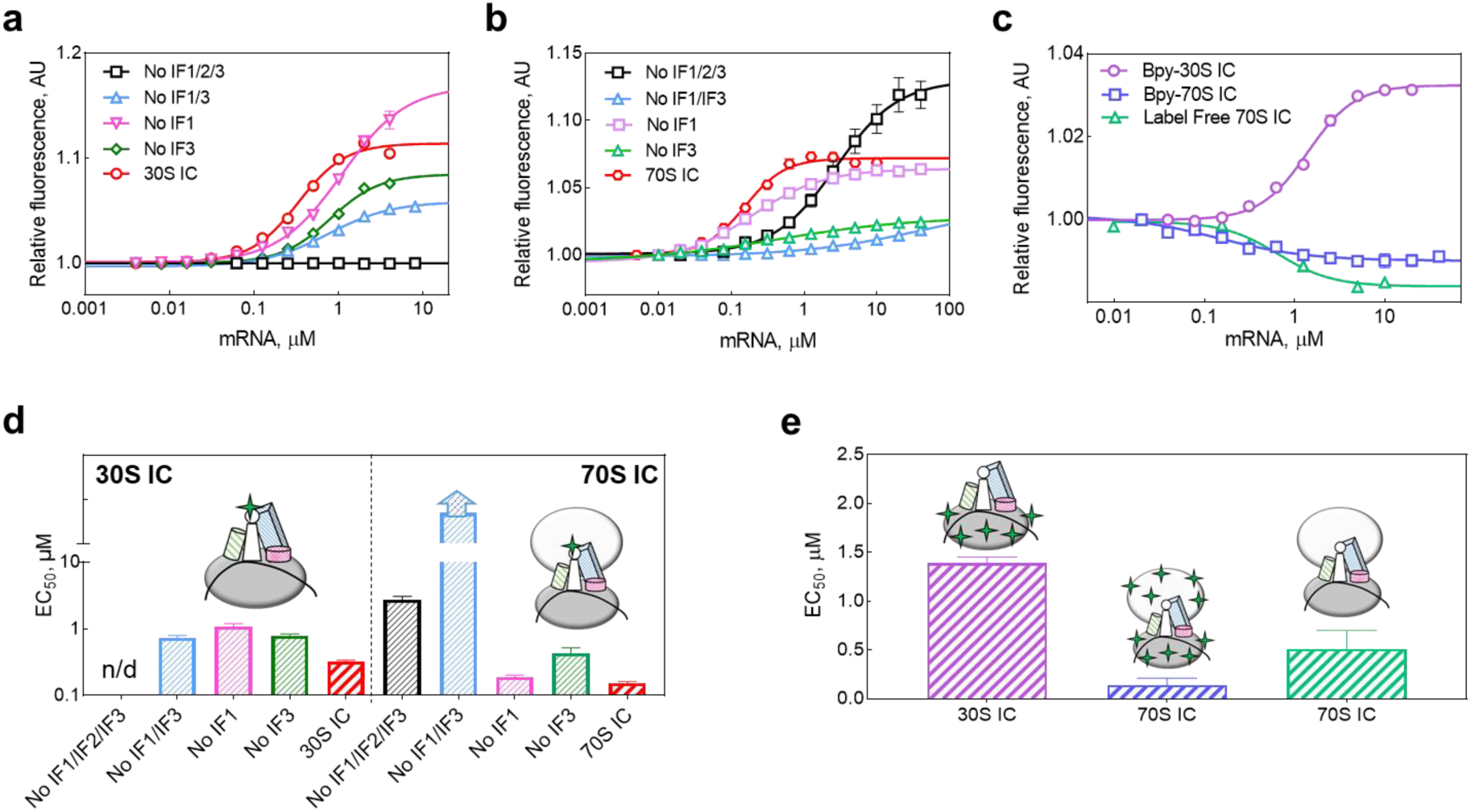
MST-analysis of stepwise translation initiation complex assembly. 30S IC and 70S IC formation as a function of mRNA concentration depending on the presence of initiation factors using Bpy-tRNA^fMet^ (a, b) or (c) Bpy-30S/Bpy-70S as reporter ligands or label-free conditions. EC_50_ values for 30S IC and 70S IC formation using Bpy-tRNA^fMet^ (d), Bpy-30S/Bpy-70S ribosomes or label-free conditions (e). Error bars indicate standard deviations from triplicates.

Binding of Bpy-tRNA^fMet^ to the 70S ribosome is characterized by a decrease in fluorescence signal with minimal change in signal amplitude (Fig. 4c). Addition of mRNA to the 70S•Bpy-tRNA^fMet^ bimolecular complex reversed the thermophoretic direction and increased signal amplitude (Fig. 5b, d). As observed for 70S IC, absence of any IFs decreased mRNA affinity relative to the complete system; exclusion of IF3 alone as reduced affinity >2-fold, while combined omission of IF3 and IF1 caused a >50-fold decrease affinity accompanied by reduced thermophoretic amplitude. The absence of IF1 had minimal impact on 70S IC assembly efficiency. We further analysis of 70S IC formation efficiency as a function of mRNA concentration using Bpy-70S ribosomes, in the presence or absence of initiation factors (Fig. S3d). The Bpy-70S•mRNA•tRNA^fMet^ ternary complex formed with comparable efficiency (3.1 ± 0.2 μM; Fig. S3d) to that measured using Bpy-tRNA^fMet^ (2.7 ± 0.3 μM; Fig. 5d). In the absence of IF1/IF3, mRNA binding to Bpy-70S•IF2•tRNA^fMet^ during 70S IC formation occurred with higher affinity (4.8 ± 0.9 μM; Fig. S3d) than for complexes using Bpy-tRNA^fMet^ as the reporter molecule (Fig. 5d).

We further used three complementary MST strategies to monitor 30S IC and 70S IC formation, employing labeled ribosomes or label-free configurations (Fig. 5c, e). Ligand affinities during IC assembly were comparable across all approaches and aligned with values obtained using Bpy-tRNA^fMet^ as reporter (Fig. 5d, e). The directionality of thermophoretic signals was identical for Bpy-tRNA^fMet^ and Bpy-30S during 30S IC formation (Fig. 5a, c). For 70S IC assembly Bpy-70S or unlabeled ligands produced descending MST traces with increasing mRNA concentration—opposite to the Bpy-tRNA^fMet^ signal (Fig. 5c)—while IFs absence experiments (lacking IF1, IF3, or all Ifs) with Bpy-70S yielded ascending traces as to Bpy-tRNA^fMet^ (Fig. S3d). All the configurations delivered consistent EC_50_ values (0.3–1 μM).

### MST Detection of Translation Inhibitors

Inhibitors of diverse chemical nature frequently target critical stages of translation, ultimately arresting or disordering protein synthesis [88–91]. These compounds bind to specific functional centers on the ribosome and disrupt key steps codon recognition, peptide bond formation and other. Such specificity minimizes impact on host eukaryotic ribosomes, making antibiotics cornerstone therapy against bacterial infections [89, 92–95]. Quantifying the binding affinities of antibiotics to their targets on the ribosome is an important parameter for developing novel therapeutics and combating emerging antibiotic resistance.

Streptomycin (Str), an aminoglycoside antibiotic, binds protein S12 and 16S rRNA within the 30S subunit and induces translational errors [96–99]. Despite extensive studies of streptomycin’s mechanism of action, direct quantitative data on its affinity for the bacterial ribosome were previously absent. Dihydrostreptomycin (DHSM), the most stable reduced analog of streptomycin (with -CHO replaced by –CH_2_OH), similarly binds to 16S rRNA of the 30S subunit and induces translational misreading. Equilibrium binding studies with *E. coli* ribosomes at 25°C and 10 mM Mg²⁺ revealed *K*_D_ values of 94 nM (70S) and 1 μM (30S), respectively [97, 100, 101]. Using MST with Bpy-labeled ribosomes, we measured streptomycin dissociation constants of 0.3 ± 0.1 μM for both 30S and 70S ribosomes in the presence of 7 mM Mg^2+^, showing comparable affinities. The thermophoretic signals were in opposite directions (Fig. 6a). The obtained affinity values fell within the range reported for dihydrostreptomycin.

**FIGURE 6.**
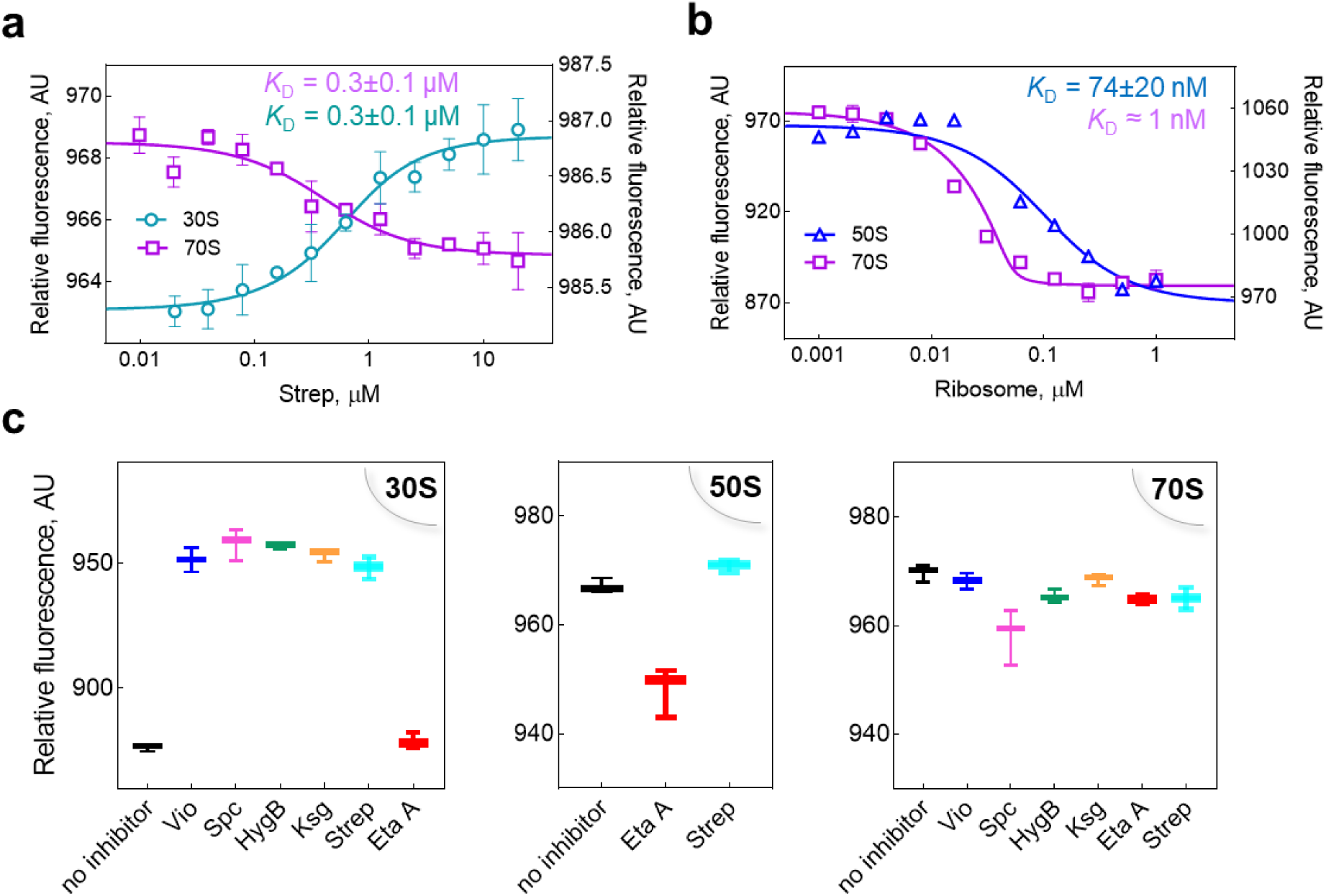
Bimolecular ribosome-inhibitor interactions by MST. (a) Streptomycin affinity for 30S/70S ribosomes: *K*_D_ = 0.3±0.1 µM (both). (b) Bpy-erythromycin affinity for 50S/70S: *K*_D_ (Bpy-Ery•50S) = 74±20 nM; *K*_D_ (Bpy-Ery•70S) ≈ 1 nM. (c) Thermophoresis shift of fluorescently labeled ribosomes in the presence of translation inhibitors: the antibiotics viomycin (Vio), spectinomycin (Spc), hygromycin B (HygB), streptomycin (Str), etamycin A (EtaA).

MST’s versatility enables the use of virtually any ligand as a fluorescent reporter. In our study, we validate this statement using macromolecules (ribosomes), medium-sized proteins (initiation factors), and small molecules (antibiotics) as reporter molecules. We employed BODIPY-labeled erythromycin (Bpy-Ery) as a reporter to probe macrolide binding with ribosome (Fig. 6b). Erythromycin reversibly binds the 50S exit tunnel through direct contacts with 23S rRNA and influences nascent peptide egress [95, 102–104]. Obtained affinities (*K*_D_ = 74±20 nM for Bpy-Ery•50S; ∼1 nM for Bpy-Ery•70S) aligned closely with prior reports [105].

In pharmaceutical development and biotechnology rapid screening of small molecules as potential inhibitors represents an indispensable strategy for discovering novel therapeutic compounds. Microscale thermophoresis enables high-throughput screening of ligand specificity with high accuracy, requiring minimal sample volumes and time. Therefore, we screened changes in the mobility of Bpy-labeled ribosomes resulting from their binding to known antibiotics targeting the 30S subunit (viomycin, spectinomycin, hygromycin B, kasugamycin, streptomycin) [88, 89, 98, 106–112] and the 50S subunit (etamycin A) [113]. Pronounced mobility shifts were observed with isolated 30S and 50S subunits in complexes with the analyzed inhibitors, whereas 70S complexes exhibited muted thermophoretic responses.

### MST Assessment of Inhibitor Effects on Translation Initiation

Elucidating the mechanisms action of translation inhibitors is essential for developing novel antimicrobials and understanding bacterial resistance profiles. Moreover, comprehensively studied inhibitors serve as probes for exploring fundamental aspects of translation. Therefore, we examined the effects of antibiotics, DNA aptamers, and a proline-rich antimicrobial peptide on 70S initiation complex formation using diverse fluorescent reporters.

We first investigated streptomycin binding to the 30S•IFs complex by monitoring IF1^Cy5^, given that both agents target the ribosomal A site. Str titration produced upward MST traces, opposite to the 30S•IF1^Cy5^ binding signal (Fig. 4a), that can be suggesting Str reduces IF1 affinity (Fig. 7a). Dose-response fitting yielded an EC_50_ of 0.14 μM.

**FIGURE 7.**
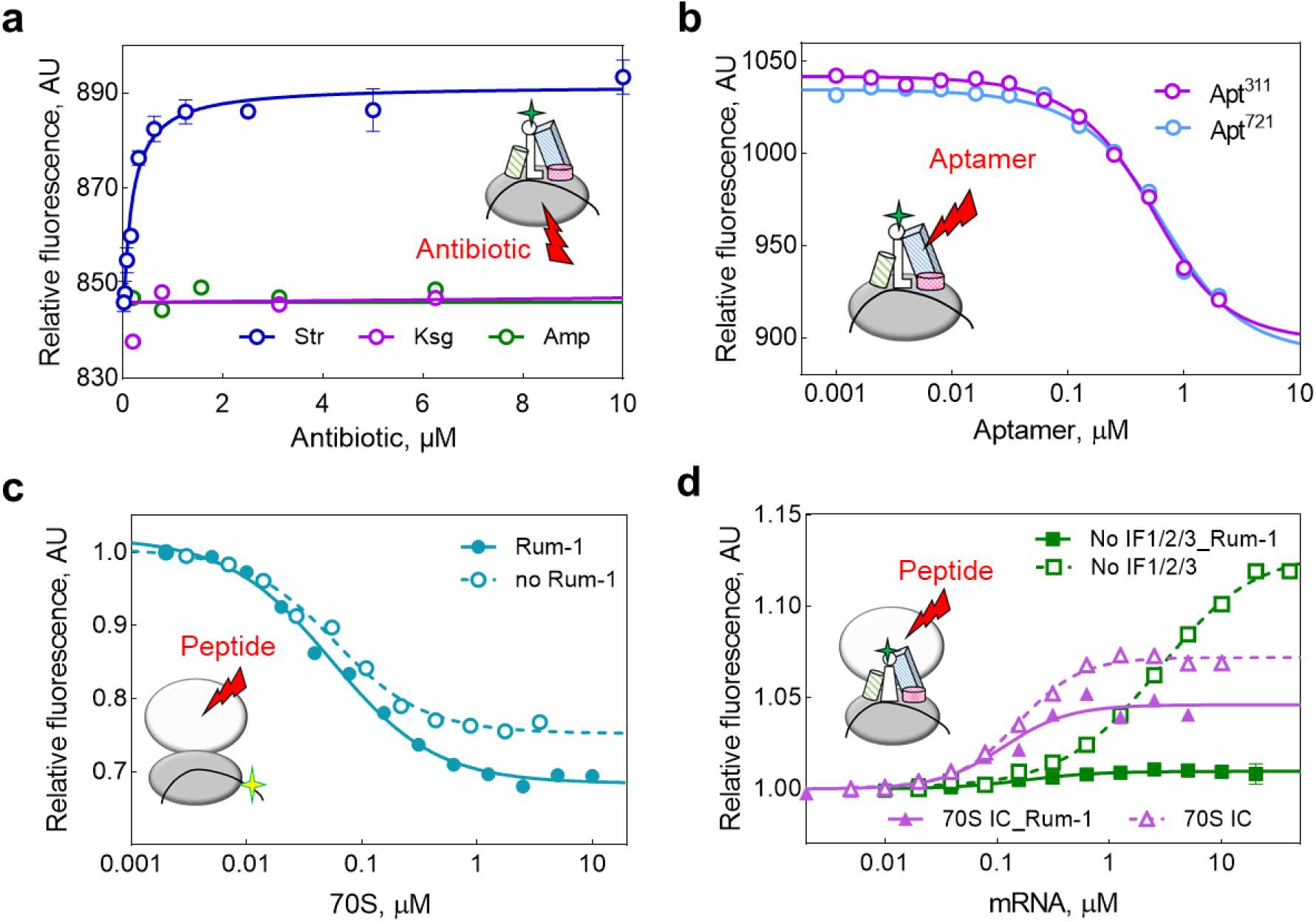
MST analysis of translation initiation complex formation in the presence of diverse inhibitors. (a) Monitoring the effects of A site binder streptomycin (Str), E site binder kasugamycin (Ksg), and translation-unrelated ampicillin (Amp) using IF1^Cy5^. (b) Effects of IF2-targeting aptamers (Apt^311^, Apt^721^) on 30S IC formation, monitored using Bpy-tRNA^fMet^. (c) Binding curves of the bimolecular 70S•mRNA-Flu complex in the presence of rumicidin-1. (d) Efficiency of 70S IC formation in the presence of the proline-rich rumicidin-family peptide Rum-1. Analysis of 70S IC formation in the presence of Rum-1, dependent on initiation factors (IFs), conducted via mRNA titration using Bpy-tRNA^fMet^ as the reporter ligand.

Another antibiotic targeting the 30S ribosome that we investigated was kasugamycin (Ksg). This aminoglycoside representative [115] binds to the 30S ribosomal subunit at the E site in the mRNA binding channel region between h24 and h28 of 16S rRNA, disrupting the transition from 30S PIC to 30S IC [116, 117]. Kasugamycin is thought to sterically hinder mRNA positioning at the E site, thereby affecting initiator tRNA^fMet^ placement at the ribosomal P site. Meanwhile, Ksg shows no significant effect on mRNA, IF1, or IF3 binding to the 30S ribosome [117]. In our experiments, kasugamycin presence in the 30S•IFs complex, monitored via IF1^Cy5^, did not alter the thermophoresis signal, consistent with existing data showing no effect on IF1.

We also examined the effect of ampicillin (Amp) on 30S•IFs complex formation, monitored via IF1^Cy5^. This β-lactam antibiotic from the aminopenicillin class inhibits bacterial cell wall peptidoglycan synthesis by binding to penicillin-binding proteins and exerts no direct effect on translation [118–119]. In our experiments, the presence of Amp exerted no effect on 30S•IFs formation (Fig. 7a).

Aptamers are chemically synthesized short oligonucleotides selected by SELEX (Systematic Evolution of Ligands by EXponential enrichment) that specifically bind target molecules [120–121]. We next employed the 30S IC assembly assay to evaluate IF-targeting DNA aptamers using Bpy-tRNA^fMet^—which binds IF2—as the reporter ligand (Fig. 7b). Titration of Apt^311^ and Apt^721^, both specific for IF2, produced downward MST traces [122], yielding EC_50_ values of 0.3 and 0.4 μM, respectively, upon dose-response fitting.

Proline-rich antimicrobial peptides represent promising antibacterial candidates, exhibiting broad-spectrum activity against bacteria, viruses, fungi, and cancer cells, and offering an alternative to conventional drugs amid rising antibiotic resistance [123–125]. Given the established finding that Rum-1 occupies the 50S exit tunnel, sterically impeding P and A site tRNA accommodation at the peptidyl transferase center and thereby inhibiting peptide bond formation [114], we examined the effects of this proline-rich rumicidin-family peptide on 70S initiation complex formation efficiency (Fig. 7c, d, S4). Rum-1 did not perturb mRNA affinity in the 70S•mRNA bimolecular complex, though a modest thermophoretic amplitude increase suggested altered complex mobility without direct competition at the mRNA binding site (Fig. 7c). However, Rum-1 enhanced mRNA affinity in 15.5-fold in the 70S•mRNA•Bpy-tRNA^fMet^ ternary complex, accompanied by a 15-fold reduction in MST amplitude. Addition of initiation factors partially counteracted this effect, modestly reducing mRNA affinity while diminishing MST amplitude 1.7-fold (Fig. 7d, S4).

## DISCUSSION

Translation initiation in prokaryotes constitutes a tightly regulated, multicomponent process that governs the efficiency and fidelity of protein synthesis (Fig. 1). Biochemical study of this pathway faces significant challenges due to the complex system of cooperative and sequential interactions between the ribosome, mRNA, initiator tRNA^fMet^, and initiation factors, in addition to limitations of traditional approaches such as radiolabeling. Biophysical methods including surface plasmon resonance (SPR) and isothermal titration calorimetry (ITC) are commonly employed to determine biomolecular affinities. We utilize a fluorescence-based analytical platform centered on microscale thermophoresis, which provides a robust, near-native environment for translation initiation studies (Fig. 2). The MST-method offers key advantages over SPR and ITC, including no requirement for sample immobilization, lower sensitivity to buffer conditions, rapid experimental throughput, and minimal sample consumption. However, microscale thermophoresis requires one interaction partner to bear a fluorescent label, which can be limiting in cases where labeling is challenging or undesirable; at the same time, this requirement imparts flexibility by allowing selection of the reporter molecule, enabling analysis across multiple labeling strategies and configurations.

In our study, we performed a sequential analysis of molecular affinities of ligands during the formation of 30S and 70S initiation complexes, employing each of them as the fluorescent reporter in turn. We introduce a rapid BODIPY-labeling protocol for ribosomes and subunits (Fig. 3a), with labeling efficiency and functional integrity—including formation of active 70S initiation complexes capable of peptide bond formation—confirmed in control tests (Figs. S1a, 3b, c, d). Maintaining conformational stability and resistance to aggregation in protein preparations is critical for preserving functional activity, particularly upon modifications; here, we comprehensively confirmed ribosomal integrity across extensive control tests—including, for the first time, nanoDSF application to ribosomes under varying ionic conditions—demonstrating no significant structural perturbations from BODIPY labeling (Figs. 3e, S1c; Table S1).

We initiated the analysis of translation initiation using our platform by examining bimolecular interactions between 30S/70S ribosomes and ligands serving as reporter molecules (IF1^Cy5^, IF2^Cy5^, IF3^Cy5^, mRNA-Flu, and Bpy-tRNA^fMet^). We observed high-affinity binding of initiation factors and mRNA to the 30S ribosome (Fig. 4a, b, c, d). Initiator tRNA^fMet^ binding was notably less efficient, consistent with its dependence on initiation factors (Fig. 4c, d).

Microscale thermophoresis offers high sensitivity and solution-phase flexibility for dissecting dynamic supramolecular assemblies across diverse biological systems [18, 126, 127]. We utilized the MST platform to investigate translation initiation as a multicomponent system using a diverse set of fluorescently labeled molecules. By progressively increasing system complexity, we demonstrated the role of magnesium ions in mRNA and tRNA^fMet^ binding to the 30S ribosome (Fig. S3a, S3b). Quantitative analysis of 30S IC formation showed the individual and synergistic contributions of IF1, IF2, and IF3, confirming IF2’s dominant role in initiator tRNA^fMet^ recruitment [10,128] while revealing IF1/IF3-mediated modulation of mRNA and tRNA^fMet^ affinities (Fig. 5, S3d). These equilibrium measurements complement kinetic insights from stopped-flow approaches and highlights unique capacity of MST to capture cooperative binding under near-native conditions. We applied our model system to analyze the formation of both 30S and 70S initiation complexes, using labeled tRNA^fMet^ as an example of a reporter ligand, along with labeled ribosomes or a completely label-free setup in our experiments (Fig. 5). Remarkably, all the approaches yielded consistent EC_50_ values (0.3–1.2 μM) despite execution across distinct project phases, instruments, locations, and operators. This underscores the system’s experimental robustness.

Translation is a frequent target for inhibitors of diverse origins. Accordingly, developing model systems that support not only the elucidation of fundamental mechanisms but also the assessment of potential inhibitors holds substantial importance—this approach enables both decoding their mechanisms of action for therapeutic development and addressing key questions in basic translation research. In addition to analyzing the affinity of interacting molecules, the model system we present can also be used for these purposes, namely evaluating the effects of potential inhibitors on the translation initiation stage. Using microscale thermophoresis, we analyzed the effects of antibiotics, DNA aptamers, and an antimicrobial peptide on translation initiation. We showed the potential of our system as a platform for rapid screening of potential inhibitors’ binding efficiency to their target, exemplified by the interaction of known antibiotics (viomycin, spectinomycin, hygromycin B, kasugamycin, streptomycin, etamycin A) with the ribosome (Fig. 6c). As an example of targeted affinity analysis for small molecules such as the antibiotics streptomycin or erythromycin with the ribosome, we performed measurements under alternative experimental configurations in which the reporter molecule was either the target (ribosome) (Fig. 6a) or the ligand (antibiotic) (Fig. 6b). As a result, we determined that streptomycin binds the 70S ribosome and the 30S ribosomal subunit with similar affinity. Analysis of Bpy-erythromycin binding to the 70S ribosome and the 50S ribosomal subunit revealed high-affinity interactions, with dissociation constants of ≈1 nM and 74 ± 20 nM, respectively, in agreement with previously reported data [105].

In addition to analyzing bimolecular interactions, we used our model system to assess the effects of inhibitors on translation initiation within a multicomponent complex. We showed that Rum-1 also affects the formation of the 70S initiation complex, in which the presence of initiation factors plays an important role in this process. These findings open a new direction for investigating how potential antimicrobial inhibitors act on the translation process. We also demonstrated the use of this model system to analyze the effects of antibiotics, using streptomycin and kasugamycin as examples, as well as IF2-targeting DNA aptamers, on the ribosomal translation initiation complex (Fig. 7a, b).

The model system we present, based on the MST platform, provides a sensitive biophysical toolkit for dissecting bacterial translation initiation by minimizing sample consumption while preserving sensitivity and physiological relevance, thereby enabling comprehensive characterization of initiation complex intermediates and surpassing fluorescence methods that require extensive manipulation or specialized instrumentation. Its versatility may extend beyond bacterial systems to eukaryotic initiation complexes, ribosome biogenesis, elongation/termination, and other supramolecular assemblies like spliceosomes, RNPs, and transcription machineries, addressing core questions in complex assembly, regulation, and stability across organisms; nanoDSF integration further enables monitoring of protein modifications, while rapid affinity screening—including for limited samples—supports drug development.

## MATERIALS AND METHODS

### Biological preparations

#### 70S ribosomes and ribosomal subunits

70S ribosomes and 30S/50S subunits were prepared as described previously [3]. Subunits were isolated from purified 70S ribosomes by sucrose gradient centrifugation (Ti-15 rotor, Beckman Coulter) under dissociating conditions in buffer TAKM_1.25_ (50 mM Tris-HCl pH 7.5, 70 mM NH_4_Cl, 30 mM KCl, 1.25 mM MgCl_2_). Ribosome concentration was determined by measuring the absorbance at 260 nm, where 1 unit corresponds to 23 pmol of 70S, 63 pmol of 30S, and 37 pmol of 50S ribosomes, respectively.

#### Translation factors purification

Initiation factors IF1, IF2, and IF3 were purified as described previously [15]. Elongation factors EF-Tu and EF-G were isolated following established protocols [129]. Mutant variants engineered for site-specific labeling—IF1 D5C, IF2γ G810C, and IF3 E166C—were expressed and purified according to published procedures [3, 129, 130].

### Fluorescence ligands

#### BODIPY-Met-tRNA^fMet^ (Bpy-tRNA^fMet^)

The fluorescent BODIPY FL SSE dye (D6140, Invitrogen, USA) was used to generate fluorescently labeled initiator Met-tRNA^fMet^, following a labeling procedure analogous to that described previously [15].

### Fluorescein-labeled mRNA (mRNA-Flu)

Synthetic mRNA mMF1 (5′-ACUAUGUUU-3′) [15] was produced by in vitro transcription for 3 h at 37°C. Reactions contained transcription buffer (40 mM Tris-HCl pH 7.5, 15 mM MgCl_2_, 2 mM spermidine, 10 mM NaCl), 10 mM DTT, 2.5 mM NTPs, 5 mM GMP, 0.01 U/ µL inorganic pyrophosphatase, 2 U/ µL T7 RNA polymerase and 5 ng/ µL DNA template. Transcripts were purified by three sequential ethanol precipitations followed by anion-exchange chromatography on a HiTrap Q column (5 ml; 17-1151-01, GE Healthcare), equilibrated with buffer A (50 mM Tris-HCl pH 7.0, 5 mM EDTA, 0.3 M NaCl). mRNA was eluted by a linear gradient of 0.3–1.5 M NaCl in buffer A, visualized by 8 M urea-PAGE and Methylene blue staining, pooled and recovered by three additional ethanol precipitations. mRNA was fluorescently labeled with fluorescein (M041030, Sigma-Aldrich) as described previously [3].

#### Fluorescent Labeling of IFs (IF1^Cy5^, IF2^Cy5^, IF3^Cy5^)

Mutant IFs (pre-incubated with 300 µM TCEP for 10 min at 37°C) were mixed with maleimide-Cy5 (GE Healthcare Life Sciences) at a 10:1 molar ratio in labelling buffer (25 mM Tris-HCl pH 7.1, 100 mM NH_4_Cl, 10% glycerol). Reactions proceeded for 2 h at room temperature in the dark with gentle mixing every 30 min, then were quenched with 6 mM β-mercaptoethanol. Labeled proteins were purified by cation-exchange chromatography on HiTrap SP HP columns (GE Healthcare Life Sciences). IF1 and IF3 were eluted by a gradient of 50–1000 mM NH_4_Cl; IF2 by a gradient of 30–750 mM NaCl. Purified proteins were dialyzed against storage buffer (25 mM Tris-HCl pH 7.1, 200 mM NH_4_Cl, 10% glycerol, 6 mM β-mercaptoethanol), aliquoted and stored at −80°C. Fractions were analyzed by SDS-PAGE (15% for IF1/IF3; 10% for IF2) with Coomassie brilliant blue staining. Protein concentrations and degree of labelling were determined using the Cy5 extinction coefficient (ε = 250,000 M⁻¹ cm⁻¹).

### BODIPY-labeled ribosomes (Bpy-70S, Bpy-50S, Bpy-30S)

Ribosomes were fluorescently labeled at free surface amino groups of ribosomal proteins using NHS-ester BODIPY FL SSE dye (Bpy, D6140, Invitrogen). 70S ribosomes (5 µM) were incubated with Bpy (1:10 molar ratio) in HAKM_7_ buffer (50 mM HEPES pH 8.7, 70 mM NH_4_Cl, 30 mM KCl, 7 mM MgCl_2_) for 1 h at 25°C in the dark. Unbound dye was removed by centrifugation through a 1.1 M sucrose cushion in TAKM_7_ buffer (50 mM Tris-HCl pH 7.5, 70 mM NH_4_Cl, 30 mM KCl, 7 mM MgCl_2_) at 259,000 × g for 2 h at 4°C (TLS-55 rotor, Beckman Coulter). Pellets were resuspended in TAKM_7_.

Fluorescently labeled 30S and 50S subunits were obtained by dissociating Bpy-70S in TAKM_1.25_ buffer (50 mM Tris-HCl pH 7.5, 70 mM NH_4_Cl, 30 mM KCl, 1.25 mM MgCl_2_) for ≥1 h at 4°C, followed by separation through 10–40% sucrose gradient (SW28 rotor, 28,000 rpm, 19 h, 4°C) (Fig. 3A). Gradients were fractionated based on absorbance profiles at 260 nm, and the fractions containing ribosomal subunits were collected and pelleted by centrifugation (SW55 rotor, 50,000 rpm, 18 h, 4°C). Labeling efficiency was verified by 10% SDS-PAGE.

### Biochemical Reactions

#### MST experiments

MST samples were prepared on ice in the dark with fixed fluorescent reporter concentrations (Bpy-tRNA^fMet^, mRNA-Flu, IFs^Cy5^, Bpy-70S, Bpy-30S) and serial 1:1 dilutions of titrants, mixed by gentle pipetting. Mixtures were incubated for 30 min at 37°C in TAKM_7_ (50 mM Tris-HCl pH 7.5, 70 mM NH_4_Cl, 30 mM KCl, 7 mM MgCl_2_) or TAKM_20_ (50 mM Tris-HCl pH 7.5, 70 mM NH_4_Cl, 30 mM KCl, 20 mM MgCl_2_), centrifuged (13,000 rpm, 5 min, 25°C) and loaded into standard treated capillaries (MO-K022) or label-free capillaries (MO-Z022) (NanoTemper Technologies GmbH).

Samples containing Bpy or Flu fluorophores were measured on a Monolith NT.115 instrument (green LED, 40% IR laser). Samples containing Cy5-labeled components were measured on a Monolith NT.LabelFree instrument (red LED, 10% IR laser). Label-free measurements employed 10% IR laser. 30S subunits were activated for 1 h at 37°C in TAKM_20_ prior to assays.

##### Bimolecular binding

Binding affinities of Cy5-labeled IFs to 30S subunits were measured at fixed IF concentrations (0.05 µM IF1^Cy5^/0.15 µM IF2^Cy5^/0.15 µM IF3^Cy5^) with varying 30S (7.8 nM–2 µM) concentrations. mRNA binding to ribosomes was assessed using 0.025 µM mRNA-Flu reporter with 30S (3.9 nM–2 µM) or 70S (2 nM–3.5 µM) ribosomes. Initiator tRNA^fMet^ binding employed 0.5 µM Bpy-tRNA^fMet^ with 30S (1 nM–6 µM) or 70S (5 nM–10 µM). All complexes were incubated for 30 min at 37°C.

Affinity of streptomycin was analyzed using 0.5 µM Bpy-30S or 0.2 µM Bpy-70S with antibiotic concentrations from 9.8 nM to 20 µM. Erythromycin binding employed varying ribosome concentrations (0.1 nM–1 µM) at fixed 0.05 µM Bpy-Ery. Diffusion shifts of the inhibitors were assessed with 0.5 µM Bpy-30S, 0.5 µM Bpy-50S, and 0.2 µM Bpy-70S in the presence of 200 µM of each antibiotic: viomycin (Vio), spectinomycin (Spc), hygromycin B (Hyg B), kasugamycin (Ksg), streptomycin (Str) and etamycin A (Eta A).

##### Multicomponent system

The effect of Mg^2+^ on mRNA and tRNA^fMet^ binding to 30S ribosomes was analyzed using 0.5 µM Bpy-tRNA^fMet^, 4 µM mRNA, 1 µM 30S, and Mg^2+^ concentrations from 3.1 to 100 mM. IF-dependent 30S initiation complex formation was assessed with 1 µM 30S, 0.5 µM Bpy-tRNA^fMet^, 4 µM mRNA, 0.5 mM GTP, and 0.3–10 µM IFi in TAKM_7_ and TAKM_20_ buffers.

30S/70S initiation complex formation was analyzed in the presence of initiation factors using 0.5 µM Bpy-tRNA^fMet^, 1 µM 30S/70S, 2 µM IF1, 1 µM IF2, 1.5 µM IF3, 0.5 mM GTP, and mRNA (4 nM–8 µM for 30S IC; 5 nM–40 µM for 70S IC), with incubation at 37°C for 30 min. Ribosome-monitored assays employed 0.5 µM Bpy-30S and 0.2 µM Bpy-70S; label-free setups used 1 µM 70S. 70S IC formation with Bpy-70S followed identical ligand conditions. 30S IC formation as a function of IF3 (1 nM–2.5 µM) was assessed with mRNA-Flu or polyU mRNA (UUU-UUU-UUU-…) under the same conditions.

##### Inhibitors action analysis

The effects of 40 µM streptomycin, 100 µM kasugamycin and 100 µM ampicillin on initiation factor binding to 30S ribosomes were analyzed by monitoring thermodiffusion changes of 0.05 µM IF1^Cy5^, with complex ligands at 0.05 µM 30S, 0.15 µM IF2, and 0.15 µM IF3 in TAKM_7_ buffer. IF2-targeted aptamer effects on 30S IC formation were assessed using 0.5 µM Bpy-tRNA^fMet^, 1 µM 30S, 2 µM IF1, 1 µM IF2, 1.5 µM IF3, 0.5 mM GTP, and 4 µM mRNA. The aptamers (Apt^311^, Apt^721^) were activated by heating at 95°C for 5 min, followed by 30 min incubation at room temperature and tested across 0.1 nM–2 µM concentrations.

The effect of rumicidin-1 on 70S•mRNA (mRNA-Flu) and 70S IC±IFs (Bpy-tRNA^fMet^) complex formation was assessed using a twofold serial dilution starting from 20 µM. Experiment on the effect of Rum-1 on the formation of the bimolecular 70S•mRNA-Flu complex was performed in the presence of 2 µM peptide. The efficiency of 70S IC formation as a function of mRNA in the presence of Rum-1 was assessed analogously to the experiments on initiation complex formation efficiency described above. The concentrations of the ligands were 0.5 µM Bpy-tRNA^fMet^, 1 µM 70S, 2 µM IF1, 1 µM IF2, µM IF3, 0.5 mM GTP, 4 µM mRNA and 2 µM rumicidin-1.

#### nanoDSF experiments

Conformational integrity and aggregation stability of fluorescently labeled Bpy-70S, Bpy-50S, and Bpy-30S were monitored by nanoDSF using a Prometheus NT.48 (NanoTemper Technologies GmbH) in TAKxMy buffer (50 mM Tris-HCl pH 7.5, 70 mM NH_4_Cl, X mM KCl, Y mM MgCl_2_). K^+^ concentration was either 30 or 360 mM; Mg^2+^ was used at 0, 1, 5, 7, 20 or 50 mM concentration.

The samples (1 µM 70S/50S/30S/Bpy-70S/Bpy-50S/Bpy-30S in a 10 µL volume) were loaded into the glass capillaries (PR-C002, NanoTemper Technologies GmbH). Optimal LED intensity (10–30%) was determined by initial fluorescence scan, followed by heating in the range of 20–95°C at 1°C/min. Melting points (Tm) and aggregation onset temperatures were calculated using PR.ThermControl software (NanoTemper Technologies GmbH).

#### Kinetic Experiments

##### Rapid kinetics

70S association and dissociation were monitored by stopped-flow light scattering (λ_ex_ = 430 nm) on an SX-20 system (Applied Photophysics, Leatherhead, UK) using native or fluorescently labeled ribosomes and IF3. Association reactions employed final concentrations of 0.15 µM Bpy-30S and 0.05 µM Bpy-50S. Dissociation of 0.1 µM 70S/Bpy-70S was initiated upon addition of 0.3 µM IF3. All the reactions were performed in TAKM_7_ buffer at 25°C.

##### Slow kinetics

30S IC formation was analyzed using 0.5 µM Bpy-30S (30S was reactivated as for MST experiments), 0.75 µM each of IF1, IF2, IF3, 0.75 µM fMet-tRNA^fMet^, 1 mM GTP, 1 mM DTT, and 2 µM mRNA in TAKM_7_ buffer. 70S IC formation employed using 0.5 µM Bpy-70S/Bpy-30S+Bpy-50S mixtures with the same IFs, fMet-tRNA^fMet^, GTP, DTT, and mRNA concentrations in TAKM_7_. The mixtures were incubated for 60 min at 37°C, and fluorescence emission was monitored at 512 nm. The efficiency of dipeptide bond formation in the resulting complexes was assessed according to the peptide synthesis protocol.

#### Peptide bond formation

Peptide bond formation efficiency was assessed using native and fluorescently labeled 70S, 30S, 50S ribosomes. 70S ICs were formed by incubating 0.4 µM ribosomes, 1.6 µM mRNA (AUG-GUU-UUC) [16], 0.6 µM each IF1, IF2, IF3, and fMet-tRNA^fMet^ with 1 mM GTP and 1 mM DTT for 1 h at 37°C. 30S subunits were activated as described above.

Ternary complexes (0.8 µM [^14^C]Val-tRNA^Val^•EF-Tu•GTP) were prepared by incubating 1.6 µM EF-Tu with 3 mM PEP, 1 mM GTP, and pyruvate kinase (1/100 vol) for 15 min at 37°C, followed by addition of 0.8 µM [^14^C]Val-tRNA^Val^ (2:1 EF-Tu:tRNA ratio) for 5 min at 37°C.

Pre-translocation complexes were formed by mixing 70S ICs with the ternary complexes for 5 min at 37°C. Dipeptide fMet-[^14^C]Val hydrolysis from peptidyl-tRNA was induced with 0.5 M KOH (1/10 vol) for 30 min at 37°C and neutralized with glacial acetic acid (1/10 vol). Peptide bond formation efficiency was quantified by reversed-phase HPLC (RP-C8 column, Waters) using a 0–65% CH_3_CN gradient in 0.1% TFA, measuring radioactive label incorporation as described previously [131].

#### Data analysis

Relative fluorescence was calculated using MO.Control software (NanoTemper Technologies GmbH) from F_hot_ and F_cold_ regions of MST traces. Binding affinities were determined by fitting thermophoresis data as a function of ligand concentration using GraphPad Prism (GraphPad, San Diego, USA). The equilibrium dissociation constants (K_D_) were obtained by fitting a quadratic binding equation (Y=Y_Min_+(Y_Max_-Y_Min_)/2*([Z+X+K_D_]-[sqrt(((Z+X+K_D_)^2)-(4*Z*X)]))), where Z is the reporter concentration and X is the total ligand concentration. EC_50_ values were obtained by nonlinear regression using the equation (Y=Y_Min_+(Y_Max_-Y_Min_)/(1+(EC_50_/X)^n), where n is the Hill slope. All graphs show individual data points with the corresponding nonlinear regression fits.

## SUPPLEMENTARY MATERIAL

Supplemental material is available for this article.

## ACKNOWLEDGMENTS

This work was supported by the Russian Science Foundation № 23-74-10088 to D.S.V. and the Concytec/Prociencia program grant PE501079419-2022 to P.M.

## CONFLICT OF INTERESTS

The authors declare no competing interests.

## Notes

### Competing Interest Statement

The authors have declared no competing interest.

## REFERENCES

1. E. Schmitt, P.-D. Coureux, A. Monestier, E. Dubiez, и Y. Mechulam, «Start Codon Recognition in Eukaryotic and Archaeal Translation Initiation: A Common Structural Core», Int. J. Mol. Sci., т. 20, вып. 4, с. 939, фев. 2019, doi: 10.3390/ijms20040939.

2. Kozak M. Initiation of translation in prokaryotes and eukaryotes. Gene. 1999;234: 187–208. doi:10.1016/s0378-1119(99)00210-3

3. Milon P, Konevega AL, Peske F, Fabbretti A, Gualerzi CO, Rodnina MV. Transient kinetics, fluorescence, and FRET in studies of initiation of translation in bacteria. Methods Enzymol. 2007;430: 1–30. doi:10.1016/S0076-6879(07)30001-3

4. Gualerzi CO, Pon CL. Initiation of mRNA translation in bacteria: structural and dynamic aspects. Cell Mol Life Sci CMLS. 2015;72: 4341–4367. doi:10.1007/s00018-015-2010-3

5. Milón P, Maracci C, Filonava L, Gualerzi CO, Rodnina MV. Real-time assembly landscape of bacterial 30S translation initiation complex. Nat Struct Mol Biol. 2012;19: 609–615. doi:10.1038/nsmb.2285

6. Duval M, Simonetti A, Caldelari I, Marzi S. Multiple ways to regulate translation initiation in bacteria: Mechanisms, regulatory circuits, dynamics. Biochimie. 2015;114: 18–29. doi:10.1016/j.biochi.2015.03.007

7. Woese CR, Kandler O, Wheelis ML. Towards a natural system of organisms: proposal for the domains Archaea, Bacteria, and Eucarya. Proc Natl Acad Sci. 1990;87: 4576–4579. doi:10.1073/pnas.87.12.4576

8. Kyrpides NC, Woese CR. Universally conserved translation initiation factors. Proc Natl Acad Sci. 1998;95: 224–228. doi:10.1073/pnas.95.1.224

9. Rodnina MV. Translation in Prokaryotes. Cold Spring Harb Perspect Biol. 2018;10: a032664. doi:10.1101/cshperspect.a032664

10. Milón P, Rodnina MV. Kinetic control of translation initiation in bacteria. Crit Rev Biochem Mol Biol. 2012;47: 334–348. doi:10.3109/10409238.2012.678284

11. Milon P, Carotti M, Konevega AL, Wintermeyer W, Rodnina MV, Gualerzi CO. The ribosome-bound initiation factor 2 recruits initiator tRNA to the 30S initiation complex. EMBO Rep. 2010;11: 312–316. doi:10.1038/embor.2010.12

12. Studer SM, Joseph S. Unfolding of mRNA secondary structure by the bacterial translation initiation complex. Mol Cell. 2006;22: 105–115. doi:10.1016/j.molcel.2006.02.014

13. Milon P, Konevega AL, Gualerzi CO, Rodnina MV. Kinetic Checkpoint at a Late Step in Translation Initiation. Mol Cell. 2008;30: 712–720. doi:10.1016/j.molcel.2008.04.014

14. Antoun A, Pavlov MY, Lovmar M, Ehrenberg M. How initiation factors tune the rate of initiation of protein synthesis in bacteria. EMBO J. 2006;25: 2539–2550. doi:10.1038/sj.emboj.7601140

15. Vinogradova DS, Zegarra V, Maksimova E, Nakamoto JA, Kasatsky P, Paleskava A, et al. How the initiating ribosome copes with ppGpp to translate mRNAs. PLOS Biol. 2020;18: e3000593. doi:10.1371/journal.pbio.3000593

16. Maksimova EM, Vinogradova DS, Osterman IA, Kasatsky PS, Nikonov OS, Milón P, et al. Multifaceted Mechanism of Amicoumacin A Inhibition of Bacterial Translation. Front Microbiol. 2021;12: 618857. doi:10.3389/fmicb.2021.618857

17. Jerabek-Willemsen M, Wienken CJ, Braun D, Baaske P, Duhr S. Molecular Interaction Studies Using Microscale Thermophoresis. ASSAY Drug Dev Technol. 2011;9: 342–353. doi:10.1089/adt.2011.0380

18. Seidel SAI, Dijkman PM, Lea WA, Van Den Bogaart G, Jerabek-Willemsen M, Lazic A, et al. Microscale thermophoresis quantifies biomolecular interactions under previously challenging conditions. Methods. 2013;59: 301–315. doi:10.1016/j.ymeth.2012.12.005

19. Spiewla T, Grab K, Depaix A, Ziemkiewicz K, Warminski M, Jemielity J, et al. An MST-based assay reveals new binding preferences of IFIT1 for canonically and noncanonically capped RNAs. RNA. 2025;31: 181–192. doi:10.1261/rna.080089.124

20. Heilek GM, Marusak R, Meares CF, Noller HF. Directed hydroxyl radical probing of 16S rRNA using Fe(II) tethered to ribosomal protein S4. Proc Natl Acad Sci. 1995;92: 1113–1116. doi:10.1073/pnas.92.4.1113

21. Odom OW, Robbins DJ, Lynch J, Dottavio-Martin D, Kramer G, Hardesty B. Distances between 3’ ends of ribosomal ribonucleic acids reassembled into Escherichia coli ribosomes. Biochemistry. 1980;19: 5947–5954. doi:10.1021/bi00567a001

22. Auerbach T, Pioletti M, Avila H, Anagnostopoulos K, Weinstein S, Franceschi F, et al. Genetic and Biochemical Manipulations of the Small Ribosomal Subunit from Thermus thermophilus HB8. J Biomol Struct Dyn. 2000;17: 617–628. doi:10.1080/07391102.2000.10506553

23. Oakes MI, Clark MW, Henderson E, Lake JA. DNA hybridization electron microscopy: ribosomal RNA nucleotides 1392-1407 are exposed in the cleft of the small subunit. Proc Natl Acad Sci. 1986;83: 275–279. doi:10.1073/pnas.83.2.275

24. Klimova M, Senyushkina T, Samatova E, Peng BZ, Pearson M, Peske F, et al. EF-G–induced ribosome sliding along the noncoding mRNA. Sci Adv. 2019;5: eaaw9049. doi:10.1126/sciadv.aaw9049

25. Sharma H, Adio S, Senyushkina T, Belardinelli R, Peske F, Rodnina MV. Kinetics of Spontaneous and EF-G-Accelerated Rotation of Ribosomal Subunits. Cell Rep. 2016;16: 2187–2196. doi:10.1016/j.celrep.2016.07.051

26. Hershey J.W.B. (1987) Protein synthesis. In Neidhardt,F.C., Ingraham,J.L., Low,K.B., Magasanik,B., Schaechter,M. and Umbarger,H.E. (eds), Escherichia coli and Salmonella typhimurium. Cellular and Molecular Biology. ASM Press, Washington, DC, pp. 613–647

27. Liiv A, Karitkina D, Maiväli U, Remme J. Analysis of the function of E. coli 23S rRNA helix-loop 69 by mutagenesis. BMC Mol Biol. 2005 Jul 29;6:18. doi: 10.1186/1471-2199-6-18. PMID: 16053518; PMCID: PMC1190176.

28. Yusupov MM, Yusupova GZ, Baucom A, Lieberman K, Earnest TN, Cate JH, Noller HF. Crystal structure of the ribosome at 5.5 A resolution. Science. 2001;292:883–896. doi: 10.1126/science.1060089.

29. Harms J, Schluenzen F, Zarivach R, Bashan A, Gat S, Agmon I, Bartels H, Franceschi F, Yonath A. High Resolution Structure of the Large RibosomalSubunit from a Mesophilic Eubacterium. Cell. 2001;107:679–688. doi: 10.1016/S0092-8674(01)00546-3.

30. d’Aquino AE, Azim T, Aleksashin NA, Hockenberry AJ, Krüger A, Jewett MC. Mutational characterization and mapping of the 70S ribosome active site. Nucleic Acids Res. 2020 Mar 18;48(5):2777–2789. doi: 10.1093/nar/gkaa001. PMID: 32009164; PMCID: PMC7049736.

31. Görisch H, Goss DJ, Parkhurst LJ. Kinetics of ribosome dissociation and subunit association studied in a light-scattering stopped-flow apparatus. Biochemistry. 1976 Dec 28;15(26):5743–53. doi: 10.1021/bi00671a010. PMID: 795460.

32. Zhang Y, Inouye M. RatA (YfjG), an Escherichia coli toxin, inhibits 70S ribosome association to block translation initiation. Mol Microbiol. 2011 Mar;79(6):1418–29. doi: 10.1111/j.1365-2958.2010.07506.x. Epub 2011 Feb 15. PMID: 21323758; PMCID: PMC3062629.

33. Wen J, Lord H, Knutson N, Wikström M. Nano differential scanning fluorimetry for comparability studies of therapeutic proteins. Anal Biochem. 2020;593: 113581. doi:10.1016/j.ab.2020.113581

34. Johnson CM. Differential scanning calorimetry as a tool for protein folding and stability. Arch Biochem Biophys. 2013;531: 100–109. doi:10.1016/j.abb.2012.09.008

35. Real-Hohn A, Groznica M, Löffler N, Blaas D, Kowalski H. nanoDSF: In vitro Label-Free Method to Monitor Picornavirus Uncoating and Test Compounds Affecting Particle Stability. Front Microbiol. 2020;11: 1442. doi:10.3389/fmicb.2020.01442

36. Vinogradova, D.S., et al. Nanobiotechnology Reports, 2025, Vol. 20, No. 6, pp. 903–915. doi: 10.1134/S2635167625601895

37. Bashan A, Yonath A. The linkage between ribosomal crystallography, metal ions, heteropolytungstates and functional flexibility. J Mol Struct. 2008 Nov 12;890(1-3):289–294. doi: 10.1016/j.molstruc.2008.03.043. PMID: 19915655; PMCID: PMC2757297.

38. Allen SH, Wong K-P. The role of magnesium and potassium ions in the molecular mechanism of ribosome assembly: Hydrodynamic, conformational, and thermal stability studies of 16 S RNA from Escherichia coli ribosomes. Arch Biochem Biophys. 1986;249: 137–147. doi:10.1016/0003-9861(86)90568-0

39. Jelenc PC, Kurland CG. Nucleoside triphosphate regeneration decreases the frequency of translation errors. Proc Natl Acad Sci U S A. 1979;76: 3174–3178. doi:10.1073/pnas.76.7.3174

40. Bartetzko A, Nierhaus KH. Mg2+/NH4+/polyamine system for polyuridine-dependent polyphenylalanine synthesis with near in vivo characteristics. Methods Enzymol. 1988;164: 650–658. doi:10.1016/s0076-6879(88)64075-4

41. Rheinberger HJ, Nierhaus KH. The ribosomal E site at low Mg2+: coordinate inactivation of ribosomal functions at Mg2+ concentrations below 10 mM and its prevention by polyamines. J Biomol Struct Dyn. 1987;5: 435–446. doi:10.1080/07391102.1987.10506403

42. Nierhaus KH. Mg2+, K+, and the Ribosome. J Bacteriol. 2014;196: 3817–3819. doi:10.1128/JB.02297-14

43. Rivalta A, Hiregange D-G, Bose T, Rajan KS, Yonath A, Zimmerman E, et al. Ribosomes: from conserved origin to functional/medical mobility and heterogeneity. Philos Trans R Soc B Biol Sci. 2025;380: 20230393. doi:10.1098/rstb.2023.0393

44. McCarthy BJ. The effects of magnesium starvation on the ribosome content of Escherichia coli. Biochim Biophys Acta BBA - Spec Sect Nucleic Acids Relat Subj. 1962;55: 880–889. doi:10.1016/0926-6550(62)90345-6

45. Gesteland RF. Unfolding of Escherichia coli ribosomes by removal of magnesium. J Mol Biol. 1966;18: 356-IN14. doi:10.1016/S0022-2836(66)80253-X

46. Gavrilova LP, Ivanov DA, Spirin AS. Studies on the structure of ribosomes. J Mol Biol. 1966;16: 473–IN28. doi:10.1016/S0022-2836(66)80186-9

47. Gordon J, Lipmann F. Role of divalent ions in poly U-directed phenylalanine polymerization. J Mol Biol. 1967;23: 23–33. doi:10.1016/S0022-2836(67)80064-0

48. Weiss RL, Kimes BW, Morris DR. Cations and ribosome structure. III. Effects on the 30S and 50S subunits of replacing bound Mg2+ by inorganic cations. Biochemistry. 1973;12: 450–456. doi:10.1021/bi00727a014

49. Stahli C, Noll H. Structural dynamics of bacterial ribosomes: VI. Denaturation of active ribosomes by Mg2+ dependent conformational transitions. Mol Gen Genet MGG. 1977;153: 159–168. doi:10.1007/BF00264731

50. Näslund PH, Hultin T. Effects of potassium deficiency on mammalian ribosomes. Biochim Biophys Acta BBA - Nucleic Acids Protein Synth. 1970;204: 237–247. doi:10.1016/0005-2787(70)90507-1

51. Rozov A, Khusainov I, El Omari K, Duman R, Mykhaylyk V, Yusupov M, et al. Importance of potassium ions for ribosome structure and function revealed by long-wavelength X-ray diffraction. Nat Commun. 2019;10: 2519. doi:10.1038/s41467-019-10409-4

52. Yu T, Jiang J, Yu Q, Li X, Zeng F. Structural Insights into the Distortion of the Ribosomal Small Subunit at Different Magnesium Concentrations. Biomolecules. 2023;13: 566. doi:10.3390/biom13030566

53. Warner BR, Fredrick K. Contribution of an alternative 16S rRNA helix to biogenesis of the 30S subunit of the ribosome. RNA N Y N. 2024;30: 770–778. doi:10.1261/rna.079960.124

54. Abeysirigunawardena SC, Woodson SA. Differential effects of ribosomal proteins and Mg2+ ions on a conformational switch during 30S ribosome 5’-domain assembly. RNA N Y N. 2015;21: 1859–1865. doi:10.1261/rna.051292.115

55. Connolly K, Culver G. Overexpression of RbfA in the absence of the KsgA checkpoint results in impaired translation initiation. Mol Microbiol. 2013;87: 968–981. doi:10.1111/mmi.12145

56. Davis JH, Williamson JR. Structure and dynamics of bacterial ribosome biogenesis. Philos Trans R Soc B Biol Sci. 2017;372: 20160181. doi:10.1098/rstb.2016.0181

57. Klein DJ, Moore PB, Steitz TA. The contribution of metal ions to the structural stability of the large ribosomal subunit. RNA. 2004;10: 1366–1379. doi:10.1261/rna.7390804

58. Nissen P, Ippolito JA, Ban N, Moore PB, Steitz TA. RNA tertiary interactions in the large ribosomal subunit: The A-minor motif. Proc Natl Acad Sci. 2001;98: 4899–4903. doi:10.1073/pnas.081082398

59. Zitomer RS, Flaks JG. Magnesium dependence and equilibrium of the Escherichia coli ribosomal subunit association. J Mol Biol. 1972;71: 263–279. doi:10.1016/0022-2836(72)90350-6

60. Uday AB, Mishra RK, Hussain T. Initiation factor 3 bound to the 30S ribosomal subunit in an initial step of translation. Proteins Struct Funct Bioinforma. 2025;93: 279–286. doi:10.1002/prot.26655

61. De Smit MH, Van Duin J. Translational Standby Sites: How Ribosomes May Deal with the Rapid Folding Kinetics of mRNA. J Mol Biol. 2003;331: 737–743. doi:10.1016/S0022-2836(03)00809-X

62. Jay G, Abrams WR, Kaempfer R. Resistance of bacterial protein synthesis to double-stranded RNA. Biochem Biophys Res Commun. 1974;60: 1357–1364. doi:10.1016/0006-291X(74)90347-7

63. Katunin VI, Semenkov YuP, Makhno VI, Kirillov SV. Comparative study of the interaction of polyuridylic acid with 30S subunits and 70S ribosomes of Escherichia coli. Nucleic Acids Res. 1980;8: 403–421. doi:10.1093/nar/8.2.403

64. Evstafieva AG, Shatsky IN, Bogdanov AA, Semenkov YP, Vasiliev VD. Localization of 5′ and 3′ ends of the ribosome-bound segment of template polynucleotides by immune electron microscopy. EMBO J. 1983;2: 799–804. doi:10.1002/j.1460-2075.1983.tb01503.x

65. Romilly C, Deindl S, Wagner EGH. The ribosomal protein S1-dependent standby site in tisB mRNA consists of a single-stranded region and a 5′ structure element. Proc Natl Acad Sci. 2019;116: 15901–15906. doi:10.1073/pnas.1904309116

66. Hüttenhofer A, Noller HF. Footprinting mRNA-ribosome complexes with chemical probes. EMBO J. 1994;13: 3892–3901. doi:10.1002/j.1460-2075.1994.tb06700.x

67. Hussain T, Llácer JL, Wimberly BT, Kieft JS, Ramakrishnan V. Large-Scale Movements of IF3 and tRNA during Bacterial Translation Initiation. Cell. 2016;167: 133–144.e13. doi:10.1016/j.cell.2016.08.074

68. Kaledhonkar S, Fu Z, Caban K, Li W, Chen B, Sun M, et al. Late steps in bacterial translation initiation visualized using time-resolved cryo-EM. Nature. 2019;570: 400–404. doi:10.1038/s41586-019-1249-5

69. Yamamoto H, Wittek D, Gupta R, Qin B, Ueda T, Krause R, et al. 70S-scanning initiation is a novel and frequent initiation mode of ribosomal translation in bacteria. Proc Natl Acad Sci. 2016;113. doi:10.1073/pnas.1524554113

70. Moll I. Translation initiation with 70S ribosomes: an alternative pathway for leaderless mRNAs. Nucleic Acids Res. 2004;32: 3354–3363. doi:10.1093/nar/gkh663

71. Leiva LE, Katz A. Regulation of Leaderless mRNA Translation in Bacteria. Microorganisms. 2022;10: 723. doi:10.3390/microorganisms10040723

72. Gualerzi CO, Brandi L, Caserta E, Garofalo C, Lammi M, La Teana A, et al. Initiation Factors in the Early Events of mRNA Translation in Bacteria. Cold Spring Harb Symp Quant Biol. 2001;66: 363–376. doi:10.1101/sqb.2001.66.363

73. Boelens R, Gualerzi CO. Structure and Function of Bacterial Initiation Factors. Curr Protein Pept Sci. 2002;3: 107–119. doi:10.2174/1389203023380765

74. Myasnikov AG, Simonetti A, Marzi S, Klaholz BP. Structure–function insights into prokaryotic and eukaryotic translation initiation. Curr Opin Struct Biol. 2009;19: 300–309. doi:10.1016/j.sbi.2009.04.010

75. Simonetti A, Marzi S, Jenner L, Myasnikov A, Romby P, Yusupova G, et al. A structural view of translation initiation in bacteria. Cell Mol Life Sci. 2009;66: 423–436. doi:10.1007/s00018-008-8416-4

76. Laursen BS, Sørensen HP, Mortensen KK, Sperling-Petersen HU. Initiation of Protein Synthesis in Bacteria. Microbiol Mol Biol Rev. 2005;69: 101–123. doi:10.1128/MMBR.69.1.101-123.2005

77. Allen GS, Frank J. Structural insights on the translation initiation complex: ghosts of a universal initiation complex. Mol Microbiol. 2007;63: 941–950. doi:10.1111/j.1365-2958.2006.05574.x

78. Julián P, Milon P, Agirrezabala X, Lasso G, Gil D, Rodnina MV, et al. The Cryo-EM Structure of a Complete 30S Translation Initiation Complex from Escherichia coli. Spahn C, editor. PLoS Biol. 2011;9: e1001095. doi:10.1371/journal.pbio.1001095

79. Goyal A, Belardinelli R, Maracci C, Milón P, Rodnina MV. Directional transition from initiation to elongation in bacterial translation. Nucleic Acids Res. 2015;43: 10700–10712. doi:10.1093/nar/gkv869

80. Yoo J, RajBhandary UL. Requirements for translation re-initiation in Escherichia coli : roles of initiator tRNA and initiation factors IF2 and IF3. Mol Microbiol. 2008;67: 1012–1026. doi:10.1111/j.1365-2958.2008.06104.x

81. Chemla Y, Peeri M, Heltberg ML, Eichler J, Jensen MH, Tuller T, et al. mRNA secondary structure stability regulates bacterial translation insulation and re-initiation. Molecular Biology; 2020. doi:10.1101/2020.02.10.941153

82. Flügel T, Schacherl M, Unbehaun A, Schroeer B, Dabrowski M, Bürger J, et al. Transient disome complex formation in native polysomes during ongoing protein synthesis captured by cryo-EM. Nat Commun. 2024;15: 1756. doi:10.1038/s41467-024-46092-3

83. Tsai Y-J, Lee H-I, Lin A. Ribosome Distribution in HeLa Cells during the Cell Cycle. Silver DL, editor. PLoS ONE. 2012;7: e32820. doi:10.1371/journal.pone.0032820

84. Bakshi S, Choi H, Weisshaar JC. The spatial biology of transcription and translation in rapidly growing Escherichia coli. Front Microbiol. 2015;6. doi:10.3389/fmicb.2015.00636

85. Simonetti A, Marzi S, Billas IML, Tsai A, Fabbretti A, Myasnikov AG, et al. Involvement of protein IF2 N domain in ribosomal subunit joining revealed from architecture and function of the full-length initiation factor. Proc Natl Acad Sci. 2013;110: 15656–15661. doi:10.1073/pnas.1309578110

86. Gualerzi C, Risuleo G, Pon CL. Initial rate kinetic analysis of the mechanism of initiation complex formation and the role of initiation factor IF-3. Biochemistry. 1977;16: 1684–1689. doi:10.1021/bi00627a025

87. Laalami S, Sacerdot C, Vachon G, Mortensen K, Sperling-Petersen HU, Cenatiempo Y, et al. Structural and functional domains of E coli initiation factor IF2. Biochimie. 1991;73: 1557–1566. doi:10.1016/0300-9084(91)90191-3

88. Wilson DN. The A–Z of bacterial translation inhibitors. Crit Rev Biochem Mol Biol. 2009;44: 393–433. doi:10.3109/10409230903307311

89. Wilson DN. Ribosome-targeting antibiotics and mechanisms of bacterial resistance. Nat Rev Microbiol. 2014;12: 35–48. doi:10.1038/nrmicro3155

90. Brandi L, Fabbretti A, La Teana A, Abbondi M, Losi D, Donadio S, et al. Specific, efficient, and selective inhibition of prokaryotic translation initiation by a novel peptide antibiotic. Proc Natl Acad Sci. 2006;103: 39–44. doi:10.1073/pnas.0507740102

91. Sutcliffe JA. Improving on nature: antibiotics that target the ribosome. Curr Opin Microbiol. 2005;8: 534–542. doi:10.1016/j.mib.2005.08.004

92. Poehlsgaard J, Douthwaite S. The bacterial ribosome as a target for antibiotics. Nat Rev Microbiol. 2005;3: 870–881. doi:10.1038/nrmicro1265

93. Tenson T, Mankin A. Antibiotics and the ribosome. Mol Microbiol. 2006;59: 1664–1677. doi:10.1111/j.1365-2958.2006.05063.x

94. Lin J, Zhou D, Steitz TA, Polikanov YS, Gagnon MG. Ribosome-Targeting Antibiotics: Modes of Action, Mechanisms of Resistance, and Implications for Drug Design. Annu Rev Biochem. 2018;87: 451–478. doi:10.1146/annurev-biochem-062917-011942

95. Krawczyk SJ, Leśniczak-Staszak M, Gowin E, Szaflarski W. Mechanistic Insights into Clinically Relevant Ribosome-Targeting Antibiotics. Biomolecules. 2024;14: 1263. doi:10.3390/biom14101263

96. Gromadski KB, Rodnina MV. Streptomycin interferes with conformational coupling between codon recognition and GTPase activation on the ribosome. Nat Struct Mol Biol. 2004;11: 316–322. doi:10.1038/nsmb742

97. Biswas DK, Gorini L. The Attachment Site of Streptomycin to the 30S Ribosomal Subunit. Proc Natl Acad Sci. 1972;69: 2141–2144. doi:10.1073/pnas.69.8.2141

98. Demirci H, Murphy F, Murphy E, Gregory ST, Dahlberg AE, Jogl G. A structural basis for streptomycin-induced misreading of the genetic code. Nat Commun. 2013;4: 1355. doi:10.1038/ncomms2346

99. Paternoga H, Crowe-McAuliffe C, Bock LV, Koller TO, Morici M, Beckert B, et al. Structural conservation of antibiotic interaction with ribosomes. Nat Struct Mol Biol. 2023;30: 1380–1392. doi:10.1038/s41594-023-01047-y

100. Chang FN, Flaks JG. Binding of dihydrostreptomycin to Escherichia coli ribosomes: characteristics and equilibrium of the reaction. Antimicrob Agents Chemother. 1972 Oct;2(4):294–307. doi: 10.1128/AAC.2.4.294. PMID: 4133236; PMCID: PMC444310.

101. Carter AP, Clemons WM, Brodersen DE, Morgan-Warren RJ, Wimberly BT, Ramakrishnan V. Functional insights from the structure of the 30S ribosomal subunit and its interactions with antibiotics. Nature. 2000;407: 340–348. doi:10.1038/35030019

102. Schlünzen F, Zarivach R, Harms J, Bashan A, Tocilj A, Albrecht R, et al. Structural basis for the interaction of antibiotics with the peptidyl transferase centre in eubacteria. Nature. 2001;413: 814–821. doi:10.1038/35101544

103. Sothiselvam S, Liu B, Han W, Ramu H, Klepacki D, Atkinson GC, et al. Macrolide antibiotics allosterically predispose the ribosome for translation arrest. Proc Natl Acad Sci U S A. 2014;111: 9804–9809. doi:10.1073/pnas.1403586111

104. Vester B, Douthwaite S. Macrolide Resistance Conferred by Base Substitutions in 23S rRNA. Antimicrob Agents Chemother. 2001;45: 1–12. doi:10.1128/AAC.45.1.1-12.2001

105. Pichkur EB, Paleskava A, Tereshchenkov AG, Kasatsky P, Komarova ES, Shiriaev DI, et al. Insights into the improved macrolide inhibitory activity from the high-resolution cryo-EM structure of dirithromycin bound to the E. coli 70S ribosome. RNA. 2020;26: 715–723. doi:10.1261/rna.073817.119

106. Stanley RE, Blaha G, Grodzicki RL, Strickler MD, Steitz TA. The structures of the anti-tuberculosis antibiotics viomycin and capreomycin bound to the 70S ribosome. Nat Struct Mol Biol. 2010;17: 289–293. doi:10.1038/nsmb.1755

107. Holm M, Borg A, Ehrenberg M, Sanyal S. Molecular mechanism of viomycin inhibition of peptide elongation in bacteria. Proc Natl Acad Sci U S A. 2016;113: 978–983. doi:10.1073/pnas.1517541113

108. Zhang L, Wang Y-H, Zhang X, Lancaster L, Zhou J, Noller HF. The structural basis for inhibition of ribosomal translocation by viomycin. Proc Natl Acad Sci. 2020;117: 10271–10277. doi:10.1073/pnas.2002888117

109. Carter SD, Hampton CM, Langlois R, Melero R, Farino ZJ, Calderon MJ, et al. Ribosome-associated vesicles: A dynamic subcompartment of the endoplasmic reticulum in secretory cells. Sci Adv. 2020;6: eaay9572. doi:10.1126/sciadv.aay9572

110. Borovinskaya MA, Shoji S, Holton JM, Fredrick K, Cate JHD. A Steric Block in Translation Caused by the Antibiotic Spectinomycin. ACS Chem Biol. 2007;2: 545–552. doi:10.1021/cb700100n

111. Borovinskaya MA, Shoji S, Fredrick K, Cate JHD. Structural basis for hygromycin B inhibition of protein biosynthesis. RNA. 2008;14: 1590–1599. doi:10.1261/rna.1076908

112. Peske F, Savelsbergh A, Katunin VI, Rodnina MV, Wintermeyer W. Conformational Changes of the Small Ribosomal Subunit During Elongation Factor G-dependent tRNA–mRNA Translocation. J Mol Biol. 2004;343: 1183–1194. doi:10.1016/j.jmb.2004.08.097

113. Hanh BTB, Kim TH, Park J-W, Lee D-G, Kim J-S, Du YE, et al. Etamycin as a Novel Mycobacterium abscessus Inhibitor. Int J Mol Sci. 2020;21: 6908. doi:10.3390/ijms21186908

114. Panteleev PV, Pichkur EB, Kruglikov RN, Paleskava A, Shulenina OV, Bolosov IA, et al. Rumicidins are a family of mammalian host-defense peptides plugging the 70S ribosome exit tunnel. Nat Commun. 2024;15: 8925. doi:10.1038/s41467-024-53309-y

115. Umezawa, H., Hamada, M., Suhara, Y., Hashimoto, T. & Ikekawa, T. Kasugamycin, a new antibiotic. Antimicrobial Agents Chemother. 5, 753–757 (1965).

116. Structural analysis of kasugamycin inhibition of translation Barbara S Schuwirth 1,5, J Michael Day 2,5, Cathy W Hau 1,5, Gary R Janssen 2, Albert E Dahlberg 3, Jamie H Doudna Cate 1,4, Antón Vila-Sanjurjo Nat Struct Mol Biol. Author manuscript; available in PMC: 2009 Feb 5. Published in final edited form as: Nat Struct Mol Biol. 2006 Sep 24;13(10):879–886. doi: 10.1038/nsmb1150

117. The translation inhibitors kasugamycin, edeine and GE81112 target distinct steps during 30S initiation complex formation Haaris A. Safdari, Martino Morici, Ana Sanchez-Castro, Andrea Dallapè, Helge Paternoga, Anna Maria Giuliodori, Attilio Fabbretti, Pohl Milón & Daniel N. Wilson Nature Communications volume 16, Article number: 2470 (2025)

118. Killing by Ampicillin and Ofloxacin Induces Overlapping Changes in Escherichia coli Transcription Profile Niilo Kaldalu 1, Rui Mei 2, Kim Lewis Antimicrob Agents Chemother. 2004 Mar;48(3):890–896. doi: 10.1128/AAC.48.3.890-896.2004

119. Ampicillin-controlled glucose metabolism manipulates the transition from tolerance to resistance in bacteria Ming Jiang 1,2,†, Yu-bin Su 1,3,†, Jin-zhou Ye 1,†, Hui Li 1,2,†, Su-fang Kuang 1, Jia-han Wu 1, Shao-hua Li 1, Xuan-xian Peng 1,2, Bo Peng Sci Adv. 2023 Mar 8;9(10):eade8582. doi: 10.1126/sciadv.ade8582

120. Tuerk C, Gold L. Systematic evolution of ligands by exponential enrichment: RNA ligands to bacteriophage T4 DNA polymerase. Science. 1990 Aug 3;249(4968):505–10. doi: 10.1126/science.2200121. PMID: 2200121.

121. Peñaranda K, Pereira N, Savva O, Petrelli D, Spurio R, Corrigan RM, Milon P. DNA aptamer AptERA 2 targets ERA from Staphylococcus aureus and limits GTP hydrolysis. Sci Rep. 2025 Aug 22;15(1):30879. doi: 10.1038/s41598-025-15180-9. PMID: 40847110; PMCID: PMC12373895.

122. Milon, P. et al. Synthetic aptamers for the detection of the biomarker IF2 wt and IF2 ctd of Escherichia coli. Patent No. PE20190573A1 (2019).

123. Tincho MB, Morris T, Meyer M, Pretorius A. Antibacterial Activity of Rationally Designed Antimicrobial Peptides. Int J Microbiol. 2020;2020: 1–9. doi:10.1155/2020/2131535

124. Huan Y, Kong Q, Mou H, Yi H. Antimicrobial Peptides: Classification, Design, Application and Research Progress in Multiple Fields. Front Microbiol. 2020;11: 582779. doi:10.3389/fmicb.2020.582779

125. Shulenina OV, Tolstyko EA, Konevega AL, Paleskava A. New Aspects of Protein Biosynthesis Inhibition by Proline-Rich Antimicrobial Peptides. Biochem Mosc. 2025;90: 1536–1552. doi:10.1134/S0006297925602394

126. Magnez R, Bailly C, Thuru X. Microscale Thermophoresis as a Tool to Study Protein Interactions and Their Implication in Human Diseases. Int J Mol Sci. 2022;23: 7672. doi:10.3390/ijms23147672

127. Seidel SAI, Wienken CJ, Geissler S, Jerabek-Willemsen M, Duhr S, Reiter A, et al. Label-Free Microscale Thermophoresis Discriminates Sites and Affinity of Protein–Ligand Binding. Angew Chem Int Ed. 2012;51: 10656–10659. doi:10.1002/anie.201204268

128. Roy B, Liu Q, Shoji S, Fredrick K. IF2 and unique features of initiator tRNAfMet help establish the translational reading frame. RNA Biol. 2018;15: 604–613. doi:10.1080/15476286.2017.1379636

129. Chulluncuy R, Espiche C, Nakamoto J, Fabbretti A, Milón P. Conformational Response of 30S-bound IF3 to A-Site Binders Streptomycin and Kanamycin. Antibiotics. 2016;5: 38. doi:10.3390/antibiotics5040038

130. Fei J, Wang J, Sternberg SH, MacDougall DD, Elvekrog MM, Pulukkunat DK, et al. A Highly Purified, Fluorescently Labeled In Vitro Translation System for Single-Molecule Studies of Protein Synthesis. Methods in Enzymology. Elsevier; 2010. pp. 221–259. doi:10.1016/S0076-6879(10)72008-5

131. Szkaradkiewicz K, Zuleeg T, Limmer S, Sprinzl M. Interaction of fMet-tRNAfMet and fMet-AMP with the C-terminal domain of Thermus thermophilus translation initiation factor 2. Eur J Biochem. 2000;267: 4290–4299. doi:10.1046/j.1432-1033.2000.01480.x

132. Szkaradkiewicz K, Zuleeg T, Limmer S, Sprinzl M. Interaction of fMet-tRNAfMet and fMet-AMP with the C-terminal domain of Thermus thermophilus translation initiation factor 2. Eur J Biochem. 2000;267: 4290–4299. doi:10.1046/j.1432-1033.2000.01480.x

133. Sanchez-Castro A, Peñaranda K, Dallapè A, Safdari HA, Nakamoto JA, Morici M, et al. Blocking IF3N delays bacterial translation initiation. Biochemistry; 2025. doi:10.1101/2025.03.14.643242

