## Supplementary Materials for "A microscale platform for the comprehensive analysis of bacterial translation initiation"

### Ion-dependent Thermal Stability of BODIPY-labeled Ribosomes (nanoDSF)

30S ribosomal subunits exhibited enhanced thermal stability with increasing $Mg^{2+}$ concentrations. Bpy-labeled 30S subunits fully preserved this $Mg^{2+}$-dependent conformational stability trend, displaying melting profiles comparable to those of native subunits. These results confirm and extend previous observations on the $Mg^{2+}$-dependent structural integrity of the 30S ribosomal subunit [52-54]. 50S subunit conformational stability displayed characteristic $Mg^{2+}$-dependent biphasic changes, increasing at moderate concentrations but declining at high levels. Bpy-labeled 50S subunits showed monotonic stability gains across the $Mg^{2+}$ range, revealing that the modification reduced their sensitivity to $Mg^{2+}$ variations. The initial thermal transition of 70S ribosomes ($Tm_1$) showed weak $Mg^{2+}$ dependence at low concentrations, likely reflecting stabilized 30S subunit denaturation within the 70S context, followed by cooperative unfolding of both subunits. Bpy-labeled 70S ribosomes mirrored this pattern but exhibited heightened sensitivity at $Mg^{2+}$ concentrations below 5 mM. Magnesium ions play a central role in ribosome assembly by stabilizing rRNA structure and coordinating intersubunit bridges (B1a–B7a), thereby promoting productive docking of the 30S and 50S subunits. Conversely, $Mg^{2+}$ deficiency leads to the accumulation of immature ribosomal particles (pre-30S and pre-50S) [55, 56]. Metal ions such as $Na^{+}$, $K^{+}$, and $Mg^{2+}$ are known to be critical for organizing the tertiary architecture of 23S rRNA, particularly within the peptidyl transferase center, where specifically coordinated cations stabilize the catalytic RNA framework [57, 58].

The concentration of potassium ions exerts a pronounced effect on ribosomal stability and activity. High KCl levels (approximately 360 mM) induce a loosening of rRNA, increasing the accessibility of protein-binding sites during ribosome assembly, but can also promote dissociation of *E. coli* ribosomes, presumably by counteracting $Mg^{2+}$-mediated stabilization [59]. Therefore, we examined the impact of 360 mM KCl on the conformational stability of native and fluorescently labeled ribosomes and their subunits in the presence of 1, 7 or 20 mM $Mg^{2+}$. Under high $K^{+}$ conditions, melting temperatures decreased for all the analyzed samples (Table S1). Fluorescently labeled ribosomes generally mirrored the ionic dependencies observed for native particles. Collectively, these observations support a model in which $K^{+}$ ions modulate conformational states during ribosome assembly: deficiency of $K^{+}$ impairs rRNA folding and reduces efficient 30S•50S association, whereas excessive $K^{+}$ elevates ionic strength and perturbs specific stabilizing contacts [55, 56].

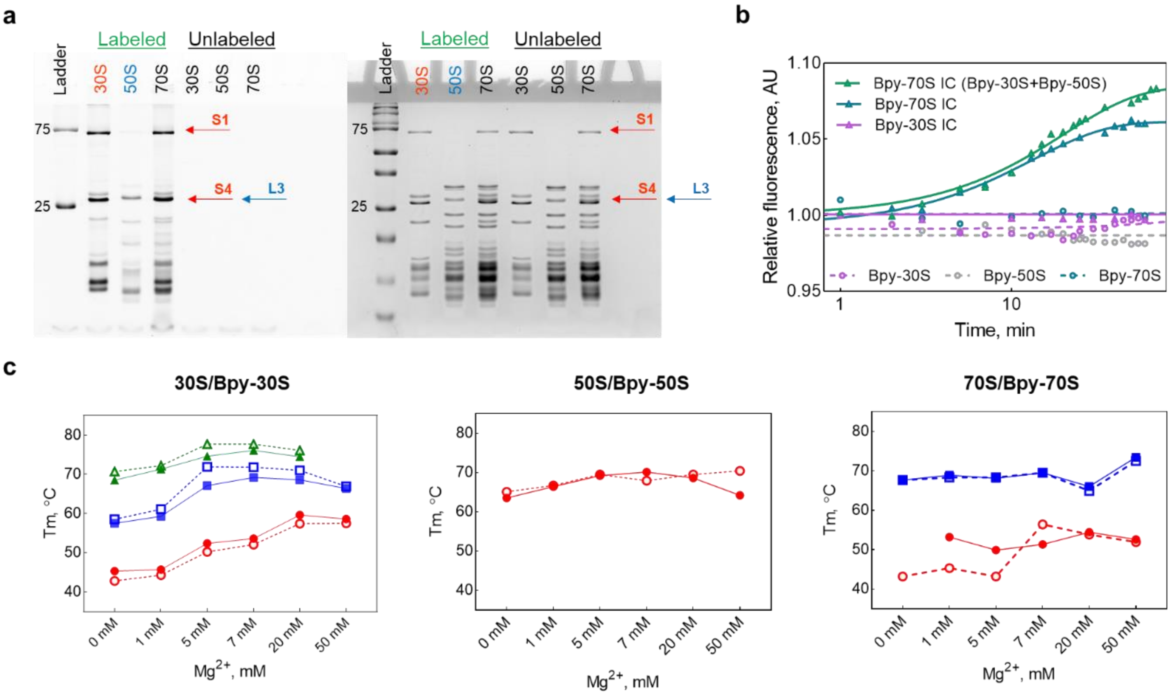


SUPPLEMENTARY FIGURE S1. Validation of the labeling efficiency and $Mg^{2+}$-dependent conformational stability of fluorescently labeled 70S ribosomes and 30S/50S ribosomal subunits. (a) 10% SDS-PAGE analysis of native and fluorescent (Bpy-labeled) 70S, 30S, 50S, showing effective labeling. (b) Kinetic analysis of 30S and 70S initiation complex formation using BODIPY-labeled 70S, 30S, and 50S ribosomal subunits. (c) Dependence of the melting temperature (Tm) values of native and fluorescently labeled 70S ribosomes and ribosomal subunits on the concentration of magnesium ions.

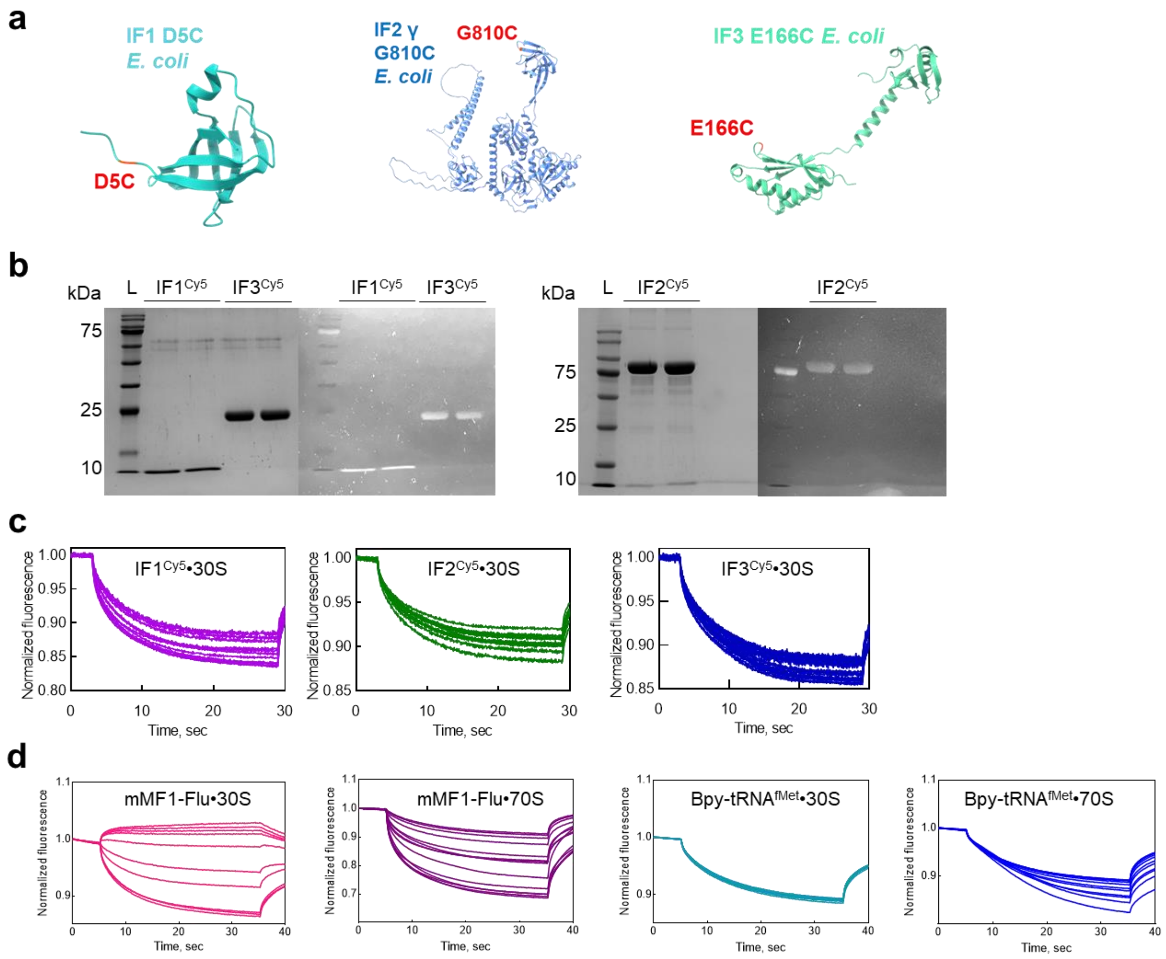


SUPPLEMENTARY FIGURE S2. Initiation factors structures for Cy5 labelling and MST time traces for bimolecular interactions between 30S/70S and IFs/mRNA/tRNA$^{fMet}$. (a) From the *E. coli* wild-type versions, mutations of the three IFs were obtained: IF1 has an exposed cysteine near the beta-barrel structure [132], the gamma isoform of IF2 has an exposed cysteine in the C-terminal domain [130], and IF3 has an exposed cysteine in the C-terminal domain [133]. The mentioned mutations are marked in red. IFs were transformed in *E. coli* BL21 (DE3) strains, expressed, purified, and labelled with maleimide Cy5 to obtain exposed cysteines. (b) 15% SDS-PAGE for IF1 and IF3 and 10% SDS-PAGE for IF2 analysis of Coomassie blue and fluorescently dye (with Cy5) of initiation factors. (c) MST traces for 30S•IF1$^{Cy5}$/IF2$^{Cy5}$/IF3$^{Cy5}$ and (d) mRNA-Flu•30S/70S and Bpy-tRNA$^{fMet}$•30S/70S interactions.

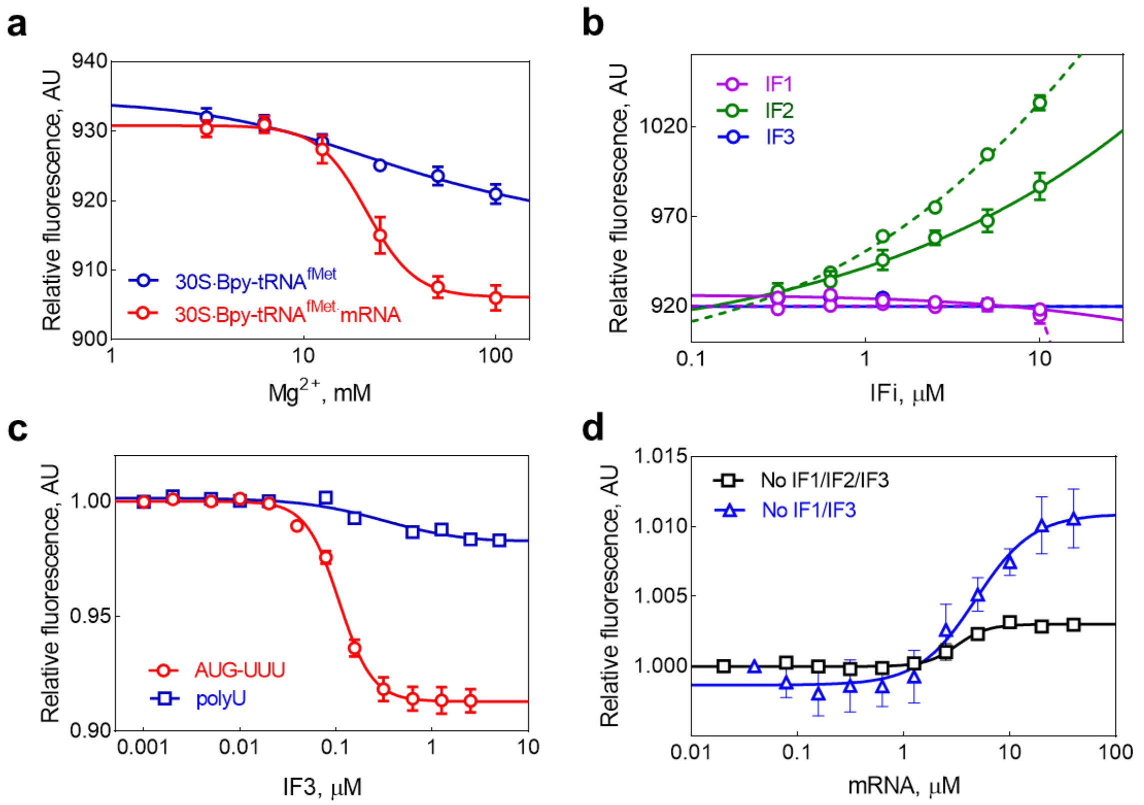


SUPPLEMENTARY FIGURE S3. Binding curves for ligand interactions with the ribosome. (a) Binding of Bpy-tRNA$^{fMet}$ to the 30S ribosomal subunit in the presence or absence of mRNA at varying $Mg^{2+}$ concentrations. (b) Stimulation of the binding of Bpy-tRNA$^{fMet}$ to mRNA and the 30S ribosome by IF2 in the presence of 7 mM (solid line) and 20 mM (dashed line) $Mg^{2+}$. (c) Efficiency of 30S initiation complex formation as a function of IF3 concentration for different mRNAs, with Bpy-tRNA$^{fMet}$ as a reporter ligand. (d) Binding of initiator tRNA$^{fMet}$ and mRNA to Bpy-labeled 70S ribosomes in the presence or absence of initiation factors.

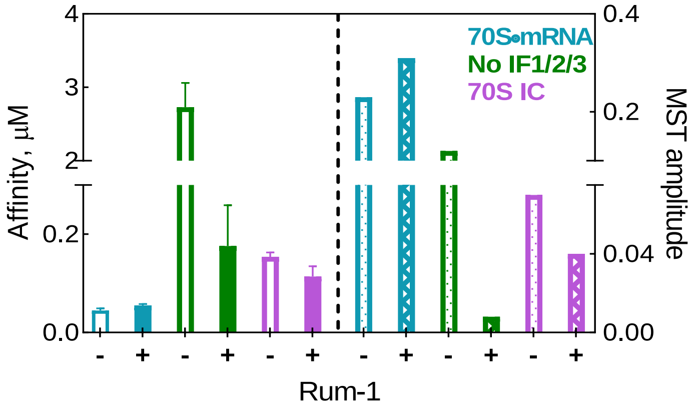


SUPPLEMENTARY FIGURE S4. Analysis of the effect of the antimicrobial peptide rumicidin-1 (Rum-1) on translation initiation. Affinity ($EC_{50}$) and MST response amplitudes for ligand binding to the 70S ribosome, shown for experiments performed in the absence and presence of Rum-1.

| | 30S | | | | | | Bpy-30S | | | | | |
|---|---|---|---|---|---|---|---|---|---|---|---|---|
| | $Tm_1$, °C | | ΔT, °C | $Tm_2$, °C | | ΔT, °C | $Tm_1$, °C | | ΔT, °C | $Tm_2$, °C | | ΔT, °C |
| $Mg^{2+}$, mM | $TAK_{30}M_i$ | $TAK_{360}M_i$ | | $TAK_{30}M_i$ | $TAK_{360}M_i$ | | $TAK_{30}M_i$ | $TAK_{360}M_i$ | | $TAK_{30}M_i$ | $TAK_{360}M_i$ | |
| 1 | 45.7 | 40.8 | ↓4.9 | 59.3 | 53.6 | ↓5.7 | 44.3 | 41.8 | ↓2.5 | 61.1 | 54.6 | ↓6.5 |
| 7 | 53.6 | 47.8 | ↓5.8 | 69.2 | 64.9 | ↓4.3 | 52.1 | 46.1 | ↓6 | 71.8 | 68.7 | ↓3.1 |
| 20 | 59.6 | – | – | 68.6 | 66.9 | ↓1.7 | 57.4 | – | – | 71.0 | 67.3 | ↓3.7 |
| | 50S | | | | | | Bpy-50S | | | | | |
| | Tm, °C | | ΔT, °C | | | | Tm, °C | | ΔT, °C | | | |
| $Mg^{2+}$, mM | $TAK_{30}M_i$ | $TAK_{360}M_i$ | | | | | $TAK_{30}M_i$ | $TAK_{360}M_i$ | | | | |
| 1 | 66.4 | (42.8) 61.5 | ↓4.9 | | | | 66.7 | (43.9) 61.6 | ↓0.4 | | | |
| 7 | 70.1 | 66.9 | ↓3.2 | | | | 67.9 | 67.5 | – | | | |
| 20 | 68.6 | 71.4 | ↑2.8 | | | | 69.5 | (52.1) 69.6 | – | | | |
| | 70S | | | | | | Bpy-70S | | | | | |
| | $Tm_1$, °C | | ΔT, °C | $Tm_2$, °C | | ΔT, °C | $Tm_1$, °C | | ΔT, °C | $Tm_2$, °C | | ΔT, °C |
| $Mg^{2+}$, mM | $TAK_{30}M_i$ | $TAK_{360}M_i$ | | $TAK_{30}M_i$ | $TAK_{360}M_i$ | | $TAK_{30}M_i$ | $TAK_{360}M_i$ | | $TAK_{30}M_i$ | $TAK_{360}M_i$ | |
| 1 | 53.2 | 48.4 | ↓4.8 | 68.9 | 66.0 | ↓2.9 | 45.3 | – | – | 68.3 | 64.8 | ↓3.5 |
| 7 | 51.3 | – | – | 69.5 | 70.9 | ↑1.4 | 56.4 | 46.3 | ↓10.1 | 69.5 | 67.6 | ↓1.9 |
| 20 | 54.4 | – | – | 66.0 | 61.3 | ↓4.7 | 53.8 | 58.6 | ↑4.8 | 64.9 | – | – |

Table S1. Melting temperatures (Tm) of native and BODIPY-labeled 70S ribosomes and of 30S and 50S ribosomal subunits at varying $Mg^{2+}$ and $K^+$ concentrations.

Vinogradova D.S.
